# Snapshots of acetyl-CoA synthesis, the final step of CO_2_ fixation in the Wood-Ljungdahl pathway

**DOI:** 10.1101/2024.08.05.606187

**Authors:** Max Dongsheng Yin, Olivier N. Lemaire, José Guadalupe Rosas Jiménez, Mélissa Belhamri, Anna Shevchenko, Gerhard Hummer, Tristan Wagner, Bonnie J. Murphy

## Abstract

In the ancient microbial Wood-Ljungdahl pathway, CO_2_ is fixed in a multi-step process ending with acetyl-CoA synthesis at the bifunctional carbon monoxide dehydrogenase/acetyl-CoA synthase complex (CODH/ACS). Here, we present catalytic snapshots of the CODH/ACS from the gas-converting acetogen *Clostridium autoethanogenum*, characterizing the molecular choreography of the overall reaction including electron transfer to the CODH for CO_2_ reduction, methyl transfer from the corrinoid iron-sulfur protein (CoFeSP) partner to the ACS active site and acetyl-CoA production. Unlike CODH, the multidomain ACS undergoes large conformational changes to form an internal connection to the CODH active site, accommodate the CoFeSP for methyl transfer and protect the reaction intermediates. Altogether, the structures allow us to draw a detailed reaction mechanism of this enzyme crucial for CO_2_ fixation in anaerobic organisms.

**One-Sentence Summary:** Structural description of key states of CO_2_ fixation by the carbon monoxide dehydrogenase/acetyl-CoA synthase complex.

## Main Text

Considered the most ancient and energy-efficient pathway for carbon dioxide (CO_2_) fixation, the Wood-Ljungdahl pathway (WLP), also known as the reductive acetyl-coenzyme A (acetyl-CoA) pathway, is responsible for 20% of annual carbon fixation (*1, 2*). The strictly anaerobic process, performed by a wide range of microbes (*e.g.*, acetogenic bacteria and methanogenic archaea) (*3*), is currently applied in gas-mitigation, biotechnologies and biofuel production (*4, 5*). The pathway consists of two branches: the methyl branch, which transforms CO_2_ into a methyl group bound to a cobalamin derivative (simplified as B12 below), and the carbonyl branch, where CO_2_ is reduced to carbon monoxide (CO). The latter reaction is catalyzed by an Fe-[Ni-3Fe-4S] cluster (C-cluster) in the carbon monoxide dehydrogenase subunit of the bifunctional carbon monoxide dehydrogenase/acetyl-CoA synthase complex (CODH/ACS) (*6–16*). The branches converge on the Ni-Ni-[4Fe-4S] cluster (A-cluster) localized in the ACS subunit (Fig. 1) to generate acetyl-CoA from CO, the methyl group, and CoA (*7, 8, 17*). Depending on the metabolism, acetyl-CoA can be converted to acetate for energy conservation, or assimilated into cell carbon (*18, 19*). Alternatively, other microbes utilize the same enzyme for the reverse process of acetyl-CoA decarbonylation (*10, 20*).

**Fig. 1.**
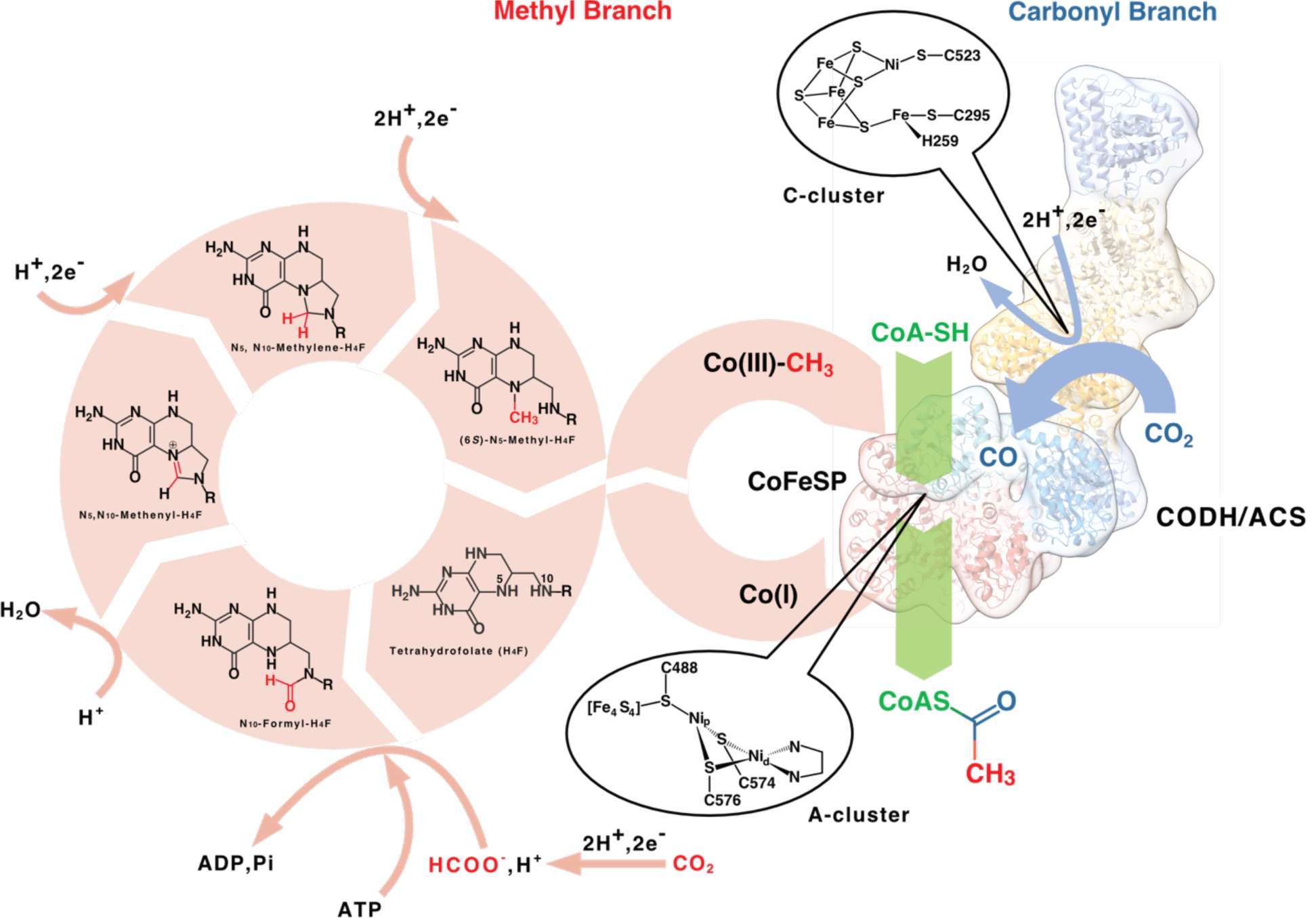
The Wood-Ljungdahl pathway. One molecule of CO_2_ is reduced in the multi-step methyl branch (pink arrows), while another is reduced at the C-cluster of CODH in the carbonyl branch (light blue arrows). The methyl and carbonyl moieties react, and the acetyl group is transferred as a thioester of coenzyme A, at the A-cluster of ACS (green arrows). For simplicity, only the bacterial WLP is depicted. The reverse decarbonylation process and the archaeal equivalent pathway are not shown. The latter depends on analogous reactions, with tetrahydromethanopterin replacing H_4_F and a formate condensation reaction being independent of ATP hydrolysis as major differences (*3*).

Structural insights into the catalytic reactions of CODH and ACS have been obtained from standalone enzymes and bifunctional CODH/ACS complexes from *Moorella thermoacetica, Carboxydothermus hydrogenoformans* and *Clostridium autoethanogenum* (*Mt, Ch and Ca*, respectively) (*7, 8, 11, 14–17, 21, 22*). However, the overall molecular mechanism of acetyl-CoA synthesis is still not fully understood due to the complexity of the reaction, which requires several additional actors and substantial structural rearrangements of the ACS.

In one of the accepted scenarios, the reaction initiates with the CO_2_ reduction at the C-cluster, which requires electron transfer from a ferredoxin (*23*). The electrons are first transferred to a solvent-exposed [4Fe-4S] cluster (D-cluster, alternatively a [2Fe-2S] cluster) (*6, 11, 24–26*) located on the symmetrical axis of the CODH dimer, before being transferred to the C-cluster through an intermediate [4Fe-4S] cluster (B-cluster). Once produced, CO is channeled to the A-cluster through a hydrophobic internal tunneling network and covalently binds as a carbonyl group to the proximal Ni (Ni_p_) (*7, 15, 21, 27*). Subsequently, the methyl-Co(III)-B12, carried by the corrinoid iron-sulfur protein (CoFeSP), interacts with the carbonylated ACS to perform methyl transfer. The methyl and carbonyl groups react to generate an acetylated A-cluster (*28*), which promotes the formation of acetyl-CoA through its reaction with the thiol group of CoA.

This reaction mechanism requires flexibility of the ACS as a prerequisite for complete turnover, as the ACS must undergo sequential reactions dependent on ferredoxin, gas trafficking, CoFeSP and CoA. The ACS is composed of three functional domains (A1, A2, and A3, from N- to C-terminus) separated by linkers that allow interdomain flexibility. Multiple conformational arrangements of the ACS have been shown through X-ray crystallography and negative-stain electron microscopy (*7, 8, 14–17, 22, 29*). However, high-resolution structures of the CODH/ACS complex with its partners or ligands are lacking.

In this study, we aimed to capture the CODH/ACS in action by visualizing the missing conformations under various protein-protein interaction or ligand-binding conditions. All presented results are derived from proteins anaerobically isolated from the biotechnologically relevant syngas converter *C. autoethanogenum*, an acetogen that we cultivate heterotrophically on fructose and H_2_/CO_2_ as reported previously (*15*). Previous studies have shown that CODH/ACS from *C. autoethanogenum* catalyzes the reversible CO oxidation with artificial electron acceptors or ferredoxin as the physiological partner, methylates the A-cluster with methyl-cobinamide (*15, 30, 31*). To structurally characterize the mechanism of acetyl-CoA synthesis, a 1:2:1 mixture of CODH/ACS heterotetramer, CoFeSP, and ferredoxin, all identified by mass spectrometry ((*15, 30*), fig. S1), was incubated with iodomethane as methyl donor and CO as carbonyl and electron donor. This mixture was plunge-frozen for cryo-EM analysis under anaerobic conditions at ∼5% CO in the gas phase. After electron microscope imaging and initial data processing, we performed three-dimensional (3D) refinement with *C_2_* symmetry, yielding maps of the rigid core of the enzyme, composed of the CODH and A1, at resolutions reaching 1.94 Å (figs. S2-3, tables S1-2). By further classifying the CODH/ACS into different states, we gain a detailed view of acetyl-CoA synthesis.

### Ferredoxin-dependent CO_2_-reduction at the C-cluster

The less well-resolved density observed at the symmetry axis of CODH was further analyzed by focused classification and local refinement without symmetry applied, resulting in a map showing a ferredoxin harboring two [4Fe-4S] clusters bound asymmetrically near the D-cluster on the *C*_2_ symmetry axis (Fig. 2A, figs. S3B, S4 and S5A). The partial occupancy of ferredoxin and its high b-factors reflect the expected transient interaction. Hydrogen bonding and hydrophobic contacts stabilize the complex (fig. S5B), forming an interaction network that could be supplemented by electrostatic attraction between positively charged residues (Lys35, Lys63) on the flexible loops of the CODH and the negatively charged area of the ferredoxin (fig. S5C). While a previous study questioned the role of the D-cluster as an electron entry/exit point due to its midpoint potential in the monofunctional CODH of *Rhodospirillum rubrum* (*23, 32*), the observed interaction and the inter-cluster distance of 8.7 Å in our structure supports ferredoxin docking and electron transfer at the D-cluster (Fig. 2A) (*6, 11, 24–26, 33*). Electrons are then transferred via the B-cluster to the catalytic C-cluster, the density of which suggests no additional ligand on the structure (fig. S6A). An initial model of the C-cluster exhibited a short (2.3 Å) distance between Ni and the pendant iron (Fe_u_), both refined with partial occupancy (∼50% and ∼60-70%, respectively). This led us to conduct mixed quantum mechanics/molecular mechanics (QM/MM) calculations using density functional theory (DFT) for the quantum region. The analysis showed that compared to the oxidized state of the C-cluster, the Ni-Fe_u_ distance decreases in the 2-electron and 4-electron reduced states C_red1(-OH)_ and C_red2(-OH)_, with the minimal distance predicted as 2.44 Å in the C_red2(-OH)_ (figs. S7-8; table S3). In a topological analysis of the electron density, the emergence of a bond critical point between Ni and Fe_u_ further indicates the formation of a metal-metal bond in the reduced states (fig. S8D-F). A shortened Ni-Fe_u_ distance was also shown in the cyanide-bound CODH crystal structure (*34*). By contrast, in the oxidized state, Ni and Fe_u_ are part of a ring formed by Ni-[Cys523]S-Fe-S. These results suggest that two electrons are stored in the metal-metal (Ni-Fe_u_) bond of the C-cluster, as previously proposed (*35*). However, we caution that the low occupancy of the metals and putative mixture of oxidation states of the C-cluster in our sample may require further structural and spectroscopic characterization. Therefore, we model the cluster with an averaged Ni-Fe_u_ distance of 2.50 Å, a still relatively short distance compared to the previous structures of the enzyme and homologues (fig. S6B-C).

**Fig. 2.**
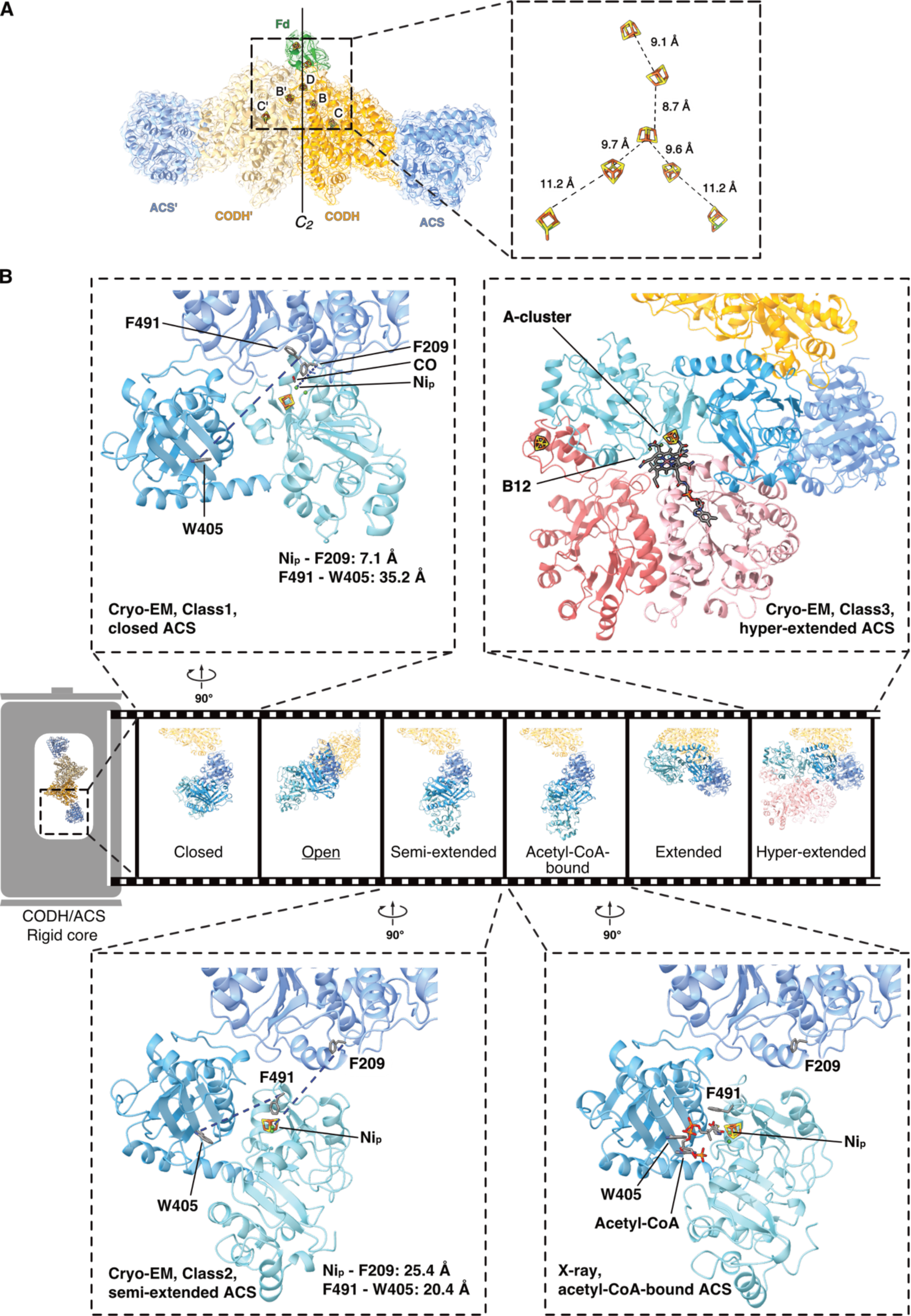
Conformational gallery of the CODH/ACS. (**A**) Ferredoxin (forest green) docks at the *C_2_* symmetry axis of CODH (gold/wheat). The symmetry axis is indicated with a vertical line. Inset: the binding conformation should allow efficient electron transfer between the [4Fe-4S] cluster of ferredoxin and the D-cluster of CODH; distances are shown as dashed lines. (**B**) The collection of flexible ACS conformations, with structures from the current study highlighted and aligned according to the A1 domain with the open state obtained from *Mt*CODH/ACS (underlined, PDB 1OAO, chain D) and the extended state (*Ca*CODH/ACS, PDB 6YTT, chain A) of ACS. The domains of ACS are shown in shades of blue, from darker at the N-terminus to lighter at the C-terminus. The Ni_p_-F209 and F491-W405 distances for the closed and semi-extended states are indicated by dashed lines, with the measured values provided alongside the structures. ACS adopts a hyper-extended state to allow binding of the methyl-donor protein CoFeSP (pink). In the hyper-extended state, the Ni_p_-F209 and F491-W405 distances are 39.8-40.5 Å and 36.8-39.4 Å, respectively, whereas in the acetyl-CoA-bound ACS, these distances are 26.7 Å and 14.9 Å, respectively (table S6). All metallocofactors are in stick representation and colored according to the element.

### Conformational spectrum of ACS allows A-cluster carbonylation

Poorly resolved density was present at the expected positions of domains A2 and A3. Using focused classification and refinement of symmetry-expanded datasets (fig. S9), we were able to separate two classes (named Class 1 and 2 in the processing workflow, refined to resolutions of 2.83 Å and 3.29 Å, respectively) with clear density for A2 and A3 but without additional density attributed to CoFeSP (figs. S3C-D, S9-10 and tables S2, S4). The rearrangement of the domains can be modeled as multi-body motion (*29*) with very low root mean square deviation (RMSD) of each domain across different conformational states and organisms (table S5). Multiple terms have been used to describe the ACS conformation in previous works, such as ‘open’, ‘closed’ and ‘extended’. Here, we introduce a reference framework for a simple comparison of these states in terms of distances between conserved sites in the three domains. In addition to the global opening/closing of the ACS due to a highly flexible loop connecting A1 to A2 (previously shown to be highly sensitive to limited proteolysis (*15*)), the A2-A3 part can potentially perform an opening-closing movement as well, allowing or occluding access to the Ni_p_ independently of the A1 position. To best characterize this dual flexibility, we propose the distances Ni_p_-F209 and F491-W405 (*Ca*ACS numbering) as proxies of interdomain 1-3 and interdomain 2-3 spacing, respectively (L_1-3_ and L_2-3,_ where ‘L’ stands for ‘length’).

The *Ca*ACS class 1 resembles the closed conformation of *Mt*CODH/ACS and *Ch*CODH/ACS (*7, 8, 14, 16*), featuring a short L_1-3_ and a long L_2-3_ (Fig. 2B, figs. S11A and S12, table S6). In the closed state, the A2-A3 space is opened and the A-cluster apposes A1 surface. A predicted CO channel emanating from the C-cluster opens into a solvent-occluded space around the A-cluster (*27*). In this state, an additional density observed on Ni_p_ is modeled as a CO bound to the tetrahedral Ni (fig. S13A-B). Well-conserved hydrophobic residues Val125, Phe209, and Phe491 are positioned similarly to their counterparts in the CO-bound state of *Mt*CODH/ACS (*14*) (fig. S13B-C). These residues were proposed to stabilize the tetrahedral geometry of the carbonylated Ni_p_, facilitate CO diffusion through internal cavities to the A-cluster, and hinder the A-cluster from adopting a methylation-compatible geometry (*14*).

By contrast, the class 2 corresponds to the semi-extended state previously described in the crystallographic structure of *Ca*CODH/ACS (PDB 6YTT, chain D) (*15*) (Fig. 2B, fig. S11B). In this state, the A3 is disengaged from the A1 and the Ni_p_ is occluded by the closing of A2-A3 (fig. S12A). The resolution is too limited to describe a bound ligand on the A-cluster (carbonyl or methyl). Class 2 likely reflects a conformation en route towards CoFeSP docking, protecting the Ni_p_ to avoid side reactions while gradually opening the interdomain A1-A3 space (table S6).

### CoFeSP interaction promotes the ACS hyper-extended state

Focused classification of the flexible ACS unveiled a third class corresponding to the CODH/ACS-CoFeSP complex (Fig. 2B, figs. S9-10, S14 and tables S2, S4). Compared to the extended state obtained by crystallography (PDB 6YTT, chain A) (*15*), ACS complexed with CoFeSP maintains a long L_1-3_ and additionally opens L_2-3_ via a 59° rotation of A3, leading to a hyper-extended state (figs. S12C and S15; table S6). Consequently, this is the sole state obtained in our study in which the A-cluster is fully accessible for methyl transfer.

The hyper-extension is maintained by three anchoring points: docking of the CoFeSP small subunit to both A1 and A3 and the interaction between the [4Fe-4S] cluster domain of the CoFeSP large subunit (1-57) and the A3 (figs. S16-17). The [4Fe-4S] cluster domain, and to a lesser extent the B12-binding domain (CoFeSP large subunit, 326-446), show the most structural differences when compared to structural homologues in complex with the activator protein or the methyltransferase (MeTr) (fig. S18) (*36, 37*). The distance between the [4Fe-4S] cluster and the A-cluster (32.3 Å, fig. S17) precludes electron transfer, agreeing with previous studies indicating no direct involvement of the CoFeSP [4Fe-4S] cluster in ACS methylation (*38, 39*). Our findings rather support the function of this domain in stabilizing the hyper-extended ACS and its interaction with CoFeSP, similar to its role during interactions with the activator protein or the MeTr (fig. S18) (*36, 37*). Residues involved in the interaction between ACS and CoFeSP are mostly conserved among bacteria and archaea (figs. S16-17), suggesting a similar mode of action across microbial kingdoms.

### Methyl transfer reaction through B12 domain motion

The hyper-extended state of the ACS enables the bulky CoFeSP to access the A-cluster (‘class 3’; Fig. 2B and Fig. 3), but additional motion of the CoFeSP is required to bring the B12 close enough for methyl transfer (*37, 40*). Via 3D variability analysis (3DVA) of the class 3, we resolve a rotational motion of the B12-binding domain (Fig. 3B). Supervised classification using three intermediate reconstructions from 3DVA led to three subsets (3A, 3B and 3C), with 3A and 3B refined to 2.71 Å and 2.65 Å, respectively (table S2). The subset 3C appeared as a mixture of two states, with the corrinoid group density stretching further towards Ni_p_ (fig. S19). By building preliminary models for each configuration and using their corresponding molmaps for supervised classification of class 3C, we obtained two substates, 3Cα and 3Cβ, refined to 2.78 Å and 2.88 Å, respectively (table S2). These maps represent snapshots of the rotational motion, during which the B12 approaches Ni_p_ (Fig. 3, movie S1), reminiscent of the motion described in the *Mt*CoFeSP complexed with its MeTr (*37*). In the 3A/3B/3Cβ sequential movement, the B12 ring progressively breaks all hydrogen bonds with the CoFeSP and establishes new ones with the ACS (Fig. 3C). These conserved hydrogen-bonding residues stabilize the B12 as it moves towards the A-cluster, with the shortest Co-Ni_p_ distance of 6.7 Å observed in 3Cβ (Fig. 3B-D), nearly sufficient for direct methyl transfer. We suppose that a transient state exists with a further shortened distance suitable for methyl transfer, as in the CoFeSP-MeTr, but this state appears too rare to be captured via classification or 3DVA. No obvious density for a methyl or carbonyl group could be detected at either the corrinoid or the Ni_p_ site. We assessed the possibility of ligand loss due to radiation damage at the site by reconstructing only early frames, corresponding to as little as 3.7 MGy (*41*), but did not observe notable differences in the density (fig. S20A).

**Fig. 3.**
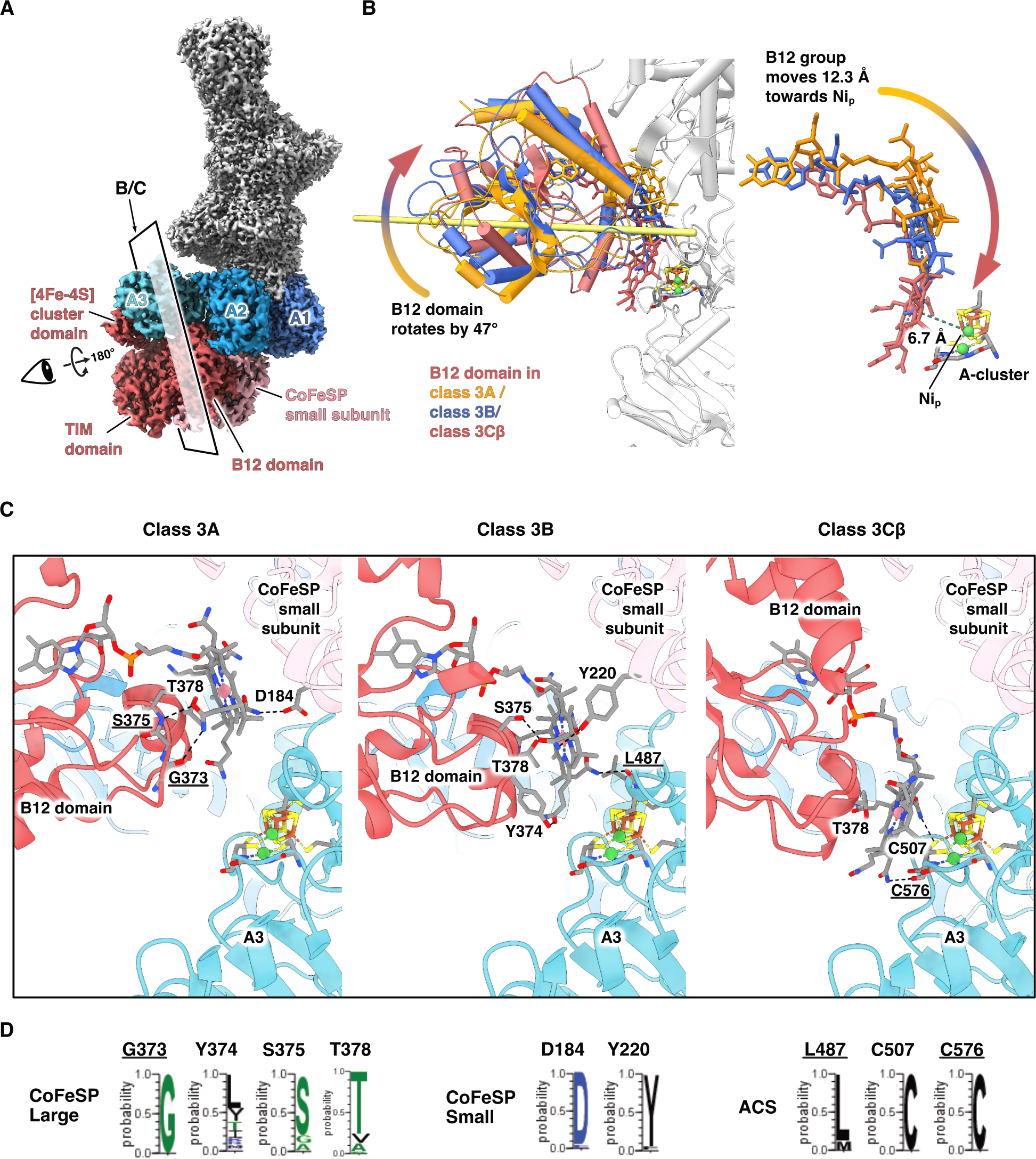
In the hyper-extended, CoFeSP-bound state, rotation of the CoFeSP B12 domain brings the B12 toward the Ni_p_. (**A**) The cryo-EM map, with the ACS in complex with a CoFeSP colored according to the established color scheme established in Fig. 2, while the rest of the complex is in gray. (**B**) The B12 domain undergoes a 47° rotation (rotation axis as a light-yellow stick) from class 3A to class 3Cβ, positioning the Co atom of B12 6.7 Å away from the Ni_p_. The B12 domain and B12 are colored according to the rotational states, and the rest of the complex is in gray. (**C**) Detailed views of the three rotational states, with key hydrogen bonds indicated by dashed lines. **(D)** Sequence conservation analysis shows that most of the B12-stabilizing residues are well conserved in bacteria and archaea. In both (C) and (D), residues involved in hydrogen-bonding via their main chain atoms are underlined.

For the states containing no external ligand at Ni_p_, our refined models consistently exhibited relatively short distances between the Ni_p_ and S1 sulfide of the [4Fe-4S] cluster (fig. S20B). To test the chemical feasibility of this configuration, we carried out a QM/MM analysis (fig. S7), using the best-resolved structure, *i.e.*, class 3B. Our calculations suggested that, in the absence of a fourth ligand bound to Ni^+^, a strong attractive interaction with the [4Fe-4S] cluster brings Ni_p_ into bonding distance (S21A; table S7). The topological analysis of the electron density reveals a bond critical point and a corresponding bond path connecting one of the sulfides in the cubane and the Ni_p_ cation (fig. S21B). A ring critical point is found between one of the Fe-S edges of the cluster and the Cys488-Ni_p_ bond, forming a 4-membered ring resembling one of the faces of the cubane. Previous experimental and theoretical studies proposed significant rearrangements between tetrahedral and square planar geometries during catalysis (*42*). Based on our calculations, it is feasible that a low-valent Ni_p_ is coordinated by a sulfide of the [4Fe-4S] cluster, generating the extended cubane structure observed in the experiment. Upon ligand binding, a four-coordinated Ni complex can form, recovering the usual geometry proposed for other reaction intermediates. Therefore, the Ni_p_ geometry modeled in our cryo-EM structures is consistent with the apparent absence of density for ligands, even though a low occupancy (∼45-59%) of the Ni_p_ site, the relatively low resolution and the putatively mixed state may hamper the accurate determination of the Ni_p_ geometry and the detection of ligands.

### The acetyl-CoA complex, a snapshot of the bound reaction product

Carbonyl and methyl groups bound at Ni_p_ combine to form an acetyl (*28*), which is subsequently transferred to the thiol group of CoA. Despite our efforts, we could not unambiguously detect additional cryo-EM map density in CODH/ACS treated with CoA or acetyl-CoA. However, by co-crystallizing *Ca*CODH/ACS with acetyl-CoA, we obtained a 2.93 Å X-ray structure of the product-bound complex (CODH/ACS_AC_, table S8). The global conformations resemble those of the as-isolated *Ca*CODH/ACS (*15*) (fig. S22). However, the previously semi-extended ACS now exhibits additional density spanning the A2 surface and reaching A3, modeled as acetyl-CoA, while the extended ACS of the asymmetric unit lacks such density, reflecting the strict conformational requirement for ligand binding (Fig. 4, fig. S23).

**Fig. 4.**
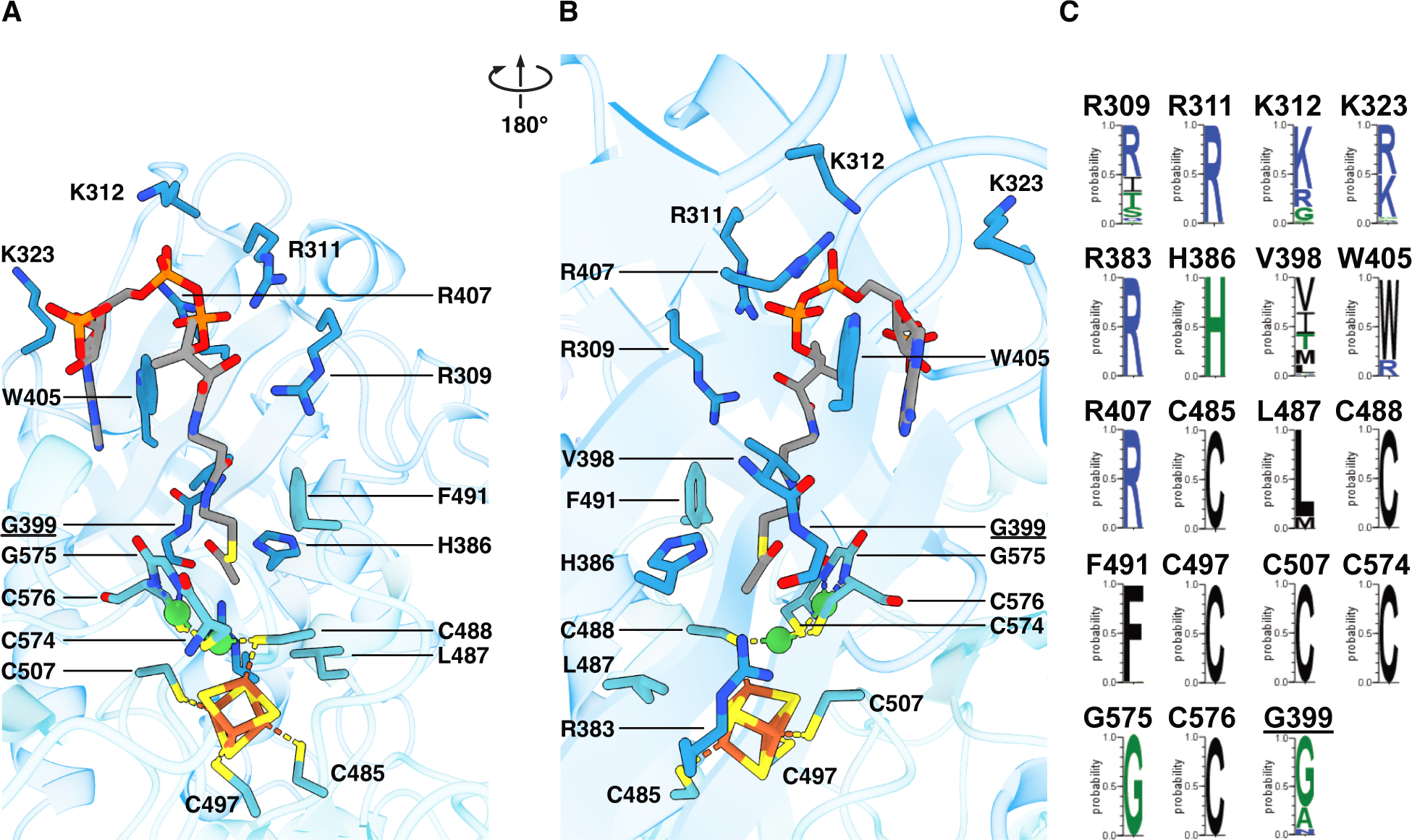
CODH/ACS in the acetyl-CoA bound state. **(A-B)** Close-up views of acetyl-CoA binding, highlighting the key residues involved in the interaction. **(C)** Conservation of the residues in bacteria and archaea. Residues involved in hydrogen-bonding via their main chain atoms are underlined.

The acetyl-CoA primarily interacts with A2 residues, with the adenine group stabilized by Trp405 via π-stacking, and the diphosphate facing a positively charged patch rich in lysines and arginines (Fig. 4, fig. S24A). The relatively high b-factors of the adenosine diphosphate moiety (fig. S24B) indicate only partial stabilization of the acetyl-CoA in the A2-A3 cleft that is restrained by the crystalline packing.

Compared to the ligand-free semi-extended ACS resolved by cryo-EM, acetyl-CoA-bound ACS undergoes a 15° rotation of A3 (fig. S25A). This further stabilizes acetyl-CoA binding via additional contacts with the A3 (Fig. 4), resolves potential clashes between A3 and acetyl-CoA (fig. S23B), and brings the A-cluster into closer proximity of the acetyl-CoA (fig. S25A). In this state, the acetyl-bearing sulfur atom is 4.13 Å away from the Ni_p_, with the acetyl group stabilized by the hydrogen-bonding with the main chain of Gly399 for the carbonyl moiety, and interaction with the His386 for the methyl moiety (Fig. 4). The flexible Phe491 swings away from the Ni_p_ (fig. S25B) to avoid steric hindrance to the CoA attack. The conserved Arg383 and His386, located near the A-cluster in the structure (Fig. 4), are likely crucial for acetyl transfer. Superposition with an acetylated A-cluster structure from a previous DFT study (*43*) indicates that Arg383 would be in the direct vicinity of the Ni_p_-bound acetyl group, putatively stabilizing it via H-bonding (fig. S25C). His386 is well positioned in our structure to facilitate deprotonation of the thiol group (Fig. 4B), consistent with previous analysis (*17*). The majority of the residues involved in acetyl-CoA stabilization in the CODH/ACS_AC_ were already suggested in *Mt*ACS (*17*), and most of these interacting residues are conserved among bacteria and archaea (Fig. 4), which suggests a strong evolutionary pressure to avoid possible clashes during the overall catalysis.

## Discussion

By providing new structural insights into the electron transfer, methyl transfer, and acetyl-CoA formation steps, our work advances decades of studies on the catalytic cycle of the CODH/ACS. The study illustrates the elegant and intricate reaction mechanism of this key anaerobic CO_2_ fixation machinery, which exhibits a wide range of ACS conformations relying on interdomain flexibility, as proposed by previous studies (*23, 29*). To simplify the nomenclature and comparison of these conformations, we introduce new metrics based on the interdomain distances L_1-3_ and L_2-3_, enabling the clustering of conformational states in a 2D landscape (Fig. 5A, table S6). The landscape generated from structurally characterized ACS allows us to construct a conformation-based model of the overall reaction of the CODH/ACS (Fig. 5B). In a sequential carbonylation-methylation scenario, the reaction would initiate in a closed state. Ferredoxin docking on the D-cluster and the subsequent electron transfer drive the CO formation, which can diffuse from the C-cluster to the A-cluster (*14*) for Ni_p_ carbonylation. The methylation step would require a hyper-extended configuration to accommodate the methyl donor CoFeSP, involving a movement of more than 37 Å of the A-cluster (fig. S26). During the movement (increasing L_1-3_), the A-cluster-carrying A3 must be undocked from the A1, which could leave the reactive carbonyl-Ni_p_ site exposed. This appears to be mitigated by closing of L_2-3_ in the semi-extended and extended states (Fig. 5A, fig. S12A-B), which shelters the A-cluster. Upon binding of CoFeSP, the hyper-extended state is stabilized, characterized by its long distance L_2-3_ and exposed A-cluster (Figs. 2B and 5A, fig. 12C and table S6). The B12 domain of CoFeSP ‘waltzes’ toward this open space, positioning the B12 close enough for the methyl transfer. After reaction of the carbonyl and methyl groups to a Ni_p_-bound acetyl group, the ACS would release the CoFeSP. Although CoA primarily interacts with A2, it must be brought near the A-cluster on A3 to further stabilize the binding and initiate acetyl transfer. Thus, acetyl-CoA formation appears to require a return to shorter L_2-3_, where A3 is rotated relative to the semi-extended state to finely tune the cleft between A2 and A3 for the nucleophilic attack of the thiol group of CoA (fig. S25A). After acetyl-CoA release, the ACS returns to the closed conformation for another catalytic cycle.

**Fig. 5.**
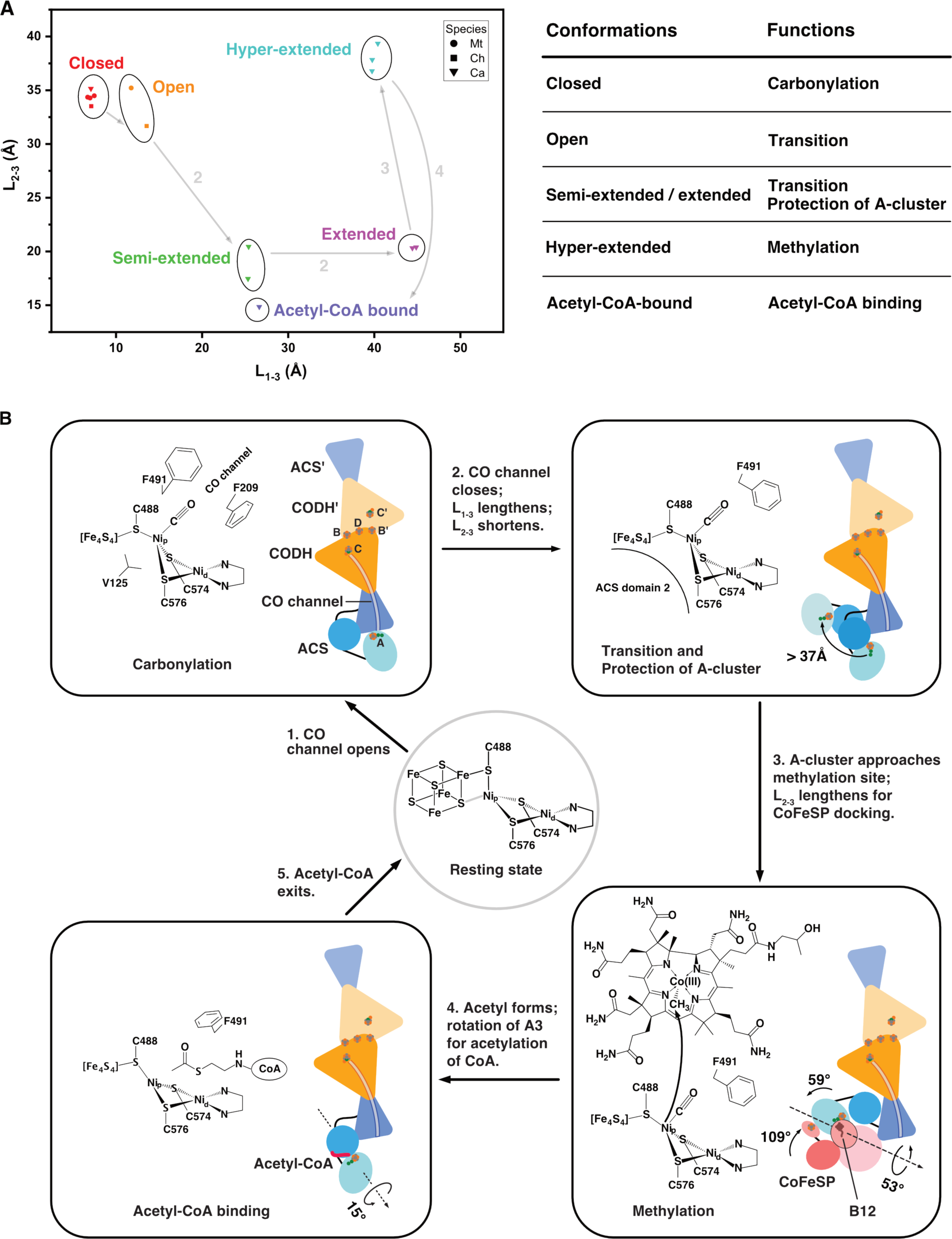
A conformation-based model of acetyl-CoA synthesis. (**A**) The conformational 2D spectrum of all known structures of CODH/ACS. One proposed reaction route, starting with carbonylation, is marked by arrows and numbered according to the model in (B). The function of each conformation is summarized in the table. (**B**) A proposed mechanism of acetyl-CoA synthesis in the sequential carbonylation-methylation scenario. Schematic representation of key CODH/ACS conformations with their functional roles, as well as the corresponding configurations of the A-cluster, are presented in the boxes. The reaction route is labeled with solid arrows, while critical domain motions during the reaction are marked by curved arrows and further illustrated in figs. S15, S18, S25A and S26. In the ligand-free resting state, the feasible Ni_p_-S coordination is indicated by a gray stick.

The proposed scenario does not exclude a reverse sequential order, *i.e.*, a sequential methylation-carbonylation process. A specific reaction order is perhaps not required, since the ACS exhibits a wide conformational spectrum in solution (Fig. 2B). We note that among all characterized *Ca*ACS structures obtained by cryo-EM from the same sample, a carbonyl ligand is observed only in the closed state, although this could also be due to experimental limitations.

The overall architecture of *Ca*CODH/ACS differs from the model *Mt*CODH/ACS or *Ch*CODH/ACS (*15*). However, aligning all reported *Ca*ACS conformation to the rigid A1 (N-terminal domain) of *Mt*ACS shows no clashes with the rigid core of *Mt*CODH. This suggests that the conformational states and reaction mechanism described here could be generalized to complexes including *Mt*CODH/ACS and *Ch*CODH/ACS, in which the N-terminal domain of ACS forms different contacts with the CODH dimer (fig. S27).

To complete the reaction landscape of ACS, several intermediate structures are still needed, including methyl-bound, CO- and methyl-bound, acetyl-bound and CoA-bound ACS. These snapshots could elucidate the mechanisms underlying the triggering or regulation of the conformational transitions, which is crucial for this reversible and central enzyme in the global carbon cycle.

## Acknowledgements

The Redox and Metalloproteins research group is grateful to the Central Electron Microscopy Facility at the Max Planck Institute of Biophysics for providing cryo-EM infrastructure and technical support. We thank Rita Zimmermann for her expertise and support in the anaerobic work. The Microbial Metabolism research group thanks the Max Planck Institute for Marine Microbiology and the Max Planck Society for their continuous support. We also thank Christina Probian and Ramona Appel for their continuous support in the Microbial Metabolism laboratory. We thank the Swiss Light Source (SLS) synchrotron, especially the staff of beamline X06DA.

## Data and materials availability

Cryo-EM structures of the CODH/ACS are available from the PDB under accession codes 9FZY (CoFeSP-bound state, class 3A), 9FZZ (CoFeSP-bound state, class 3B), 9G00 (CoFeSP-bound state, class 3Cβ), 9G01 (closed and CO-bound state), 9G02 (semi-extended state), and 9G03 (ferredoxin-bound state). Maps and half-maps are available from the EMDB under accession codes 50897 to 50909. The crystal structure of acetyl-CoA-bound CODH/ACS is available from the PDB under accession code 9G7I. Atomic coordinates from the QM/MM calculations are archived in Zenodo (*44*). The sequence alignment files used for the construction of figures are also archived in Zenodo (*45*).

## Supplementary Materials

Materials and Methods

Figs. S1 to S27

Tables S1 to S8

References (*46–81*)

Movie S1

