## Supplementary materials for "Snapshots of acetyl-CoA synthesis, the final step of CO_2_ fixation in the Wood-Ljungdahl pathway"

### Materials and Methods

#### Protein Purification

Proteins were natively extracted and purified from heterotrophically grown *Clostridium autoethanogenum* as previously described (*1, 2*). The purification protocol of CODH/ACS and ferredoxin were previously described (*1, 2*). Both were freshly prepared before structure determination.

The CoFeSP was purified multiple times through a reproducible protocol. Cell lysis and preparation of extracts were performed in an anaerobic chamber filled with an $N_2/CO_2$ atmosphere (90:10%) at room temperature. The cells were lysed via three rounds of French Press at around 1,000 PSI (6.895 MPa). To guarantee minimum oxygen contamination, the French press cell was prior flushed with $N_2$ and washed twice with an anoxic buffer. Soluble extracts were prepared by ultracentrifugation at 185,500*g* for 1 h at 4 °C. Protein purification was carried out under anaerobic conditions in a Coy tent with a $N_2/H_2$ atmosphere (97:3%) at 20 °C and under yellow light. Samples were passed through 0.2 µm filters prior to loading on chromatography columns. During purification, multi-wavelength absorbance monitoring (at 280, 415 and 550 nm) and sodium dodecyl sulfate polyacrylamide gel electrophoresis (SDS-PAGE) were used to follow the enzyme.

About 16 g (wet weight) of frozen cells were thawed and diluted with 14 ml of 50 mM Tris/HCl pH 8.0 and 2 mM dithiothreitol (DTT) before lysis. After lysis the soluble extract was diluted with 130 ml of the same buffer. The sample was loaded on 3 x 5 mL HiTrap™ DEAE Sepharose FF (GE Healthcare, Munich, Germany) equilibrated with the same buffer. After a 5 column volume (CV) washing, proteins were eluted with a 0 to 0.4 M NaCl linear gradient for 13 CV at a 2 ml/min flow rate. The CoFeSP eluted between 0.10 M and 0.14 M NaCl. Two volumes of 50 mM Tris/HCl pH 8.0 and 2 mM DTT were added to the pooled fractions before loading on a 5 ml HiTrap™ Q-Sepharose High-Performance column (GE Healthcare, Munich, Germany) column equilibrated with 25 mM Tris/HCl pH 7.6 and 2 mM DTT. After a 5 CV washing with the same buffer, proteins were eluted with a 0.05 to 0.30 M NaCl linear gradient for 12 CV at a 1 ml/min flow rate, the protein of interest eluting in a broad peak between 0.22 and 0.27 M NaCl. After fraction concentration on 10-kDa cut-off centrifugal concentrator (nitrocellulose, Vivaspin from Sartorius), contaminants were separated by size exclusion chromatography on a Superdex 200 Increase 10/300 GL (GE Healthcare, Munich, Germany) in 25 mM Tris/HCl pH 7.6, 10% (v/v) glycerol and 2 mM DTT with a flow rate of 0.4 ml/min. As initial purifications indicated a lack of absorbance for B12 and an aberrant elution profile on size exclusion chromatography, 0.1 mM Methyl-B12 was added to the protein before chromatography, and the protein eluted in a single peak. The pooled fraction was concentrated, and the protein was directly used.

#### Mass spectrometry (MS) analysis

Proteins were in-gel digested with trypsin (Promega, Germany). The resulting peptide mixtures were analyzed by liquid chromatography tandem mass spectrometry (LC-MS/MS) on a RSLCnano system UltiMate 3000 series interfaced on-line to a Velos Orbitrap LTQ hybrid mass spectrometer; the nano-LC system was equipped with Acclam PepMap 100 75 µm x 2 cm trapping column and 50 cm µPAC analytical column (all Thermo Fischer Scientific, Bremen). Peptides were separated using 75 min linear gradient, with solvent A being 0.1% aqueous formic acid, and solvent B being 0.1% formic acid in acetonitrile. Data were acquired in DDA mode using Top20 method, and precursor m/z range was 350-1,600; resolutions were 60,000 and 15,000 for precursor and fragments, respectively; dynamic exclusion time was set to 15 s. Acquired spectra were matched by Mascot software (v.2.2.04, Matrix Science, UK) with 5 ppm and 0.5 Da tolerance for precursors and fragments against *C. autoethanogenum* protein sequences in NCBI database (November 2021). The results were evaluated by Scaffold software (v.5.0.1, Proteome Software, US) using 99% and 95% protein and peptide probability thresholds, and FDR (False Discovery Rate, Scaffold option) calculated <1%.

Specimen preparation for cryo-electron microscopy

All grids for the cryo-EM study were prepared in an anaerobic chamber (Coy Laboratories) filled with 3-5% $H_2$ in $N_2$. UltrAuFoil gold grids (0.6/1, 300 mesh, Quantifoil Micro Tools GmbH) were glow-discharged (PELCO easiGlow; 90s, 15mA, 0.38 mBar air) immediately before being transferred into the chamber. Prior to grid preparation, equimolar amounts of CODH/ACS heterotetramer and ferredoxin were mixed in buffer A (25 mM Tris pH 7.6 and 2 mM DTT), gassed with CO and sealed in high performance liquid chromatography (HPLC) vials (Macherey-Nagel GmbH). CoFeSP was incubated with 1 mM methylcobalamin, exchanged into buffer A and concentrated using Amicon Ultra-0.5 centrifugal filters (Merck Millipore). To investigate CODH/ACS in complex with CoFeSP and ferredoxin, concentrated CoFeSP was incubated with iodomethane in an HPLC vial, before being injected into vials containing the CO-gassed CODH/ACS-ferredoxin mixture, using a gas-tight syringe (Hamilton company). To reduce sample interaction with the air-water interface, each sample was supplemented with 1.5 mM (0.5x CMC) fluorinated fos-choline-8 (Anatrace Products LLC), immediately before application onto the grids. Final concentrations of the CODH/ACS heterotetramer, CoFeSP, ferredoxin and iodomethane were 18 µM, 37 µM, 18 µM and 8 mM, respectively. The entire grid preparation procedure was conducted under red light due to the photosensitivity of methylcobalamin. 3 µl of sample solution was applied to each grid, which was then blotted for 6-8s with filter paper (595; GE HealthCare) and plunge-frozen into liquid ethane, using a Vitrobot Mark IV (Thermo Fisher Scientific) at 19 °C and 90% humidity. To maintain a ~5% CO atmosphere around the grids before plunge freezing, 180 ml of CO was injected into the Vitrobot chamber immediately before beginning the freezing session and this was replenished by the addition of 60 ml CO after freezing each additional two grids.

Cryo-EM imaging and data processing

Specimens were imaged by cryogenic transmission electron microscopy, using automated fast acquisition implemented via aberration-free image shift (AFIS) within EPU (Thermo Fisher Scientific) (table S1). For specimens containing CODH/ACS, CoFeSP and ferredoxin, data were acquired on a Krios G4 operating at 300 kV, equipped with a cold field emission gun, a Selectris X imaging filter and a Falcon 4 detector, with a pixel size of 0.73 Å. A total of three datasets, comprising 39,159 movies, were collected in EER format, with a dose of 70 $e^-/Å^2$ and a defocus range from -0.8 µm to -2.5 µm.

Key steps of cryo-EM data processing are illustrated in figures S2, S4 and S9. Three datasets were processed separately until the completion of 3D focused classification. Initially, beam-induced motion correction and dose weighting were carried out using MotionCor2 (*3*), implemented in RELION4 (*4*). Following this, CTF parameters were estimated for the corrected micrographs using CTFFIND 4.1.13 (*5*). Particles were initially picked using the blob picker, followed by 2D classification, *ab-initio* reconstruction and heterogeneous refinement within cryoSPARC (*6*). The coordinates of particles belonging to selected classes were then exported from CryoSPARC to RELION4 using UCSF pyem (*7*), and employed to train a TOPAZ convolutional neural network (*8*). Particles picked by TOPAZ were extracted in RELION and imported into cryoSPARC for 2D classification, *ab-initio* reconstruction and heterogeneous refinement. Following these steps, particles were selected for subsequent non-uniform refinement with 649,146 particles for the first dataset, 517,409 for the second and 706,694 for the third. Each non-uniform refinement was carried out with $C_2$ symmetry applied, incorporating per-particle defocus refinement and per-exposure-group CTF refinement on the fly. Refined particles were polished in RELION4, followed by another non-uniform refinement in cryoSPARC, which yielded consensus maps with global resolutions of 1.94 Å, 2.29 Å and 2.19 Å for the three datasets, respectively. Within each consensus map, CODH and A1 were clearly resolved, whereas density for A2 and A3 appeared blurred. To characterize the heterogeneity of ACS, particles were symmetry expanded ($C_2$) and a focused 3D classification in RELION4 was used to separate conformational states. Multiple rounds of 3D classification unveiled three conformational states of ACS: closed, semi-extended and hyper-extended states. Particle subsets corresponding to each state across datasets 1-3 were then grouped and further processed as follows.

Closed state: the corresponding subset of particles (class 1) was subjected to local refinement in cryoSPARC with a mask including the entire ACS, which yielded a focused map at a resolution of 2.83 Å (table S2).

Semi-extended state: the corresponding particles (class 2) underwent another round of focused 3D classification in RELION4, with a mask including only the flexible A2 and A3. One well-resolved class was locally refined to a resolution of 3.29 Å (table S2).

Hyper-extended state: the initial local refinement produced a focused map showing not only the ACS in a hyper-extended state but also additional density due to bound CoFeSP. Notably, the B12 domain of the CoFeSP large subunit exhibits blurred density, indicative of conformational flexibility. To explore this further, three-dimensional variability analysis (3DVA) from the cryoSPARC package was carried out on dataset 1, using a mask encompassing the A2, A3 and CoFeSP. Component 1 of the variability was due to rotation of the B12 domain (fig. S19). Three volumes were extracted from frames 3, 5 and 7 of the 3D variability display job in intermediates mode, representing the onset, midpoint and near-completion of the motion trajectory. These volumes served as reference maps for classifying particles from three datasets in cryoSPARC, focusing on the B12 domain, resulting in classes 3A, 3B and 3C, which are refined locally to resolutions of 2.71 Å, 2.65 Å and 2.65 Å, respectively (table S2). Upon close inspection of the local density around B12 in class 3C, residual motion of the corrinoid ring was observed (fig. S19). Models of B12 in two different positions were manually built into the density and used to generate molmaps in ChimeraX at a resolution of 6 Å. These molmaps, lowpass-filtered to 8 Å, were employed as reference maps for further 3D focused classification in cryoSPARC, yielding classes 3Cα and 3Cβ, which were refined locally to resolutions of 2.78 Å and 2.88 Å, respectively (table S2).

For each state described above, local resolution was estimated in cryoSPARC, and a composite map was generated with phenix.combine_focused_maps (*9*), by integrating the individual focused map with the consensus map from dataset 1, which is best resolved among those from three datasets.

Ferredoxin-bound state: additional density adjacent to the D-cluster and along the $C_2$ axis of CODH was consistently observed across all consensus maps. Focused 3D classification with a mask encompassing this region was carried out independently on each of the three datasets. Each classification yielded two classes with additional density consistent with a 2[4Fe-4S] ferredoxin from *C. autoethanogenum* (2). However, the ferredoxins in these two classes show opposite orientations about the $C_2$ symmetry axis of the CODH dimer. To combine the classes, the Euler angle describing particle rotation around the Z-axis (denoted as rlnAngleRot in RELION) was inverted by 180° in one class using the awk utility, and the resulting particle subset was merged with the other ferredoxin-containing subset. The merged subsets from three datasets were combined and refined locally with $C_1$ symmetry to a resolution of 2.1 Å (fig. S4, table S2). Local resolution was estimated within cryoSPARC (fig. S3B), and the refined map was locally filtered according to local resolution.

To assess the effect of radiation damage, early frame reconstruction was performed. During the combination step of the Bayesian polishing, only the first movie frame was used, corresponding to an electron fluence of $1e^-/Å^2$. Without further refinement of particle positions and orientations, the resulting images were used to reconstruct low-dose maps utilizing the relion_reconstruct command in RELION4 (*4*), followed by post-processing in RELION4 (*4*).

Cryo-EM model building and refinements

The initial model of the CODH/ACS rigid core comprising CODH and A1 was built by rigid body fitting of the published model (PDB ID 6YTT (*1*)) into the map. Initial models for A2, A3 and CoFeSP were generated using ModelAngelo (*10*) and subsequently manually refined in Coot (*11*). Particular focus was placed on the flexible B12 domain and the [4Fe-4S] cluster domain of CoFeSP, and models were manually adjusted while comparing with structures predicted by AlphaFold2 (*12*). Cofactors were added manually using Coot, and ligand restraints were generated via eLBOW and ReadySet within PHENIX (version 1.20.1-4487) (*9*). Iterative rounds of real space refinement and manual refinement of the models were carried out with PHENIX (*13*) and Coot. During the real space refinements, minimization_global and local_grid_search were activated. In the final round of refinement, b-factors of the whole model, occupancies of the C-cluster in all states, occupancies of the A-cluster in the CoFeSP-bound states were refined using Servalcat (*14, 15*). Figures and movies illustrating models and maps were prepared using ChimeraX (*16*).

Crystallization, data collection and structural analysis

Crystallization was performed anaerobically by initial screening at 20 °C using the sitting drop method on 96-Well MRC 2-Drop polystyrene Crystallization Plates (SWISSCI) in a Coy tent containing an $N_2/H_2$ (97:3%) atmosphere. The reservoir chamber was filled with 90 µl of crystallization condition (Ethoxylate Polymer-Based 96-Well Screen (*17*)), and the crystallization drop was formed by spotting 0.6 µl of purified protein with 0.6 µl of precipitant. The protein was crystallized at 15 mg/ml in a solution containing 25% (w/v) glycerol ethoxylate 1,000, 100 mM Tris/HCl pH 8.5 and 100 mM calcium chlorate. Before crystallization, 2 mM acetyl-CoA was added to the protein solution. The crystals were soaked in the crystallization solution supplemented with 15% (v/v) ethylene glycol for a few seconds before freezing in liquid nitrogen.

The diffraction experiments were performed at 100 K on the beamline PXIII (X06DA) from the Swiss Light Source (SLS). The data were processed and scaled with autoPROC (*18*). The data presented anisotropy (along the following axes: a = 3.227 Å, b = 3.327 Å, and c = 2.867 Å) and were further processed with STARANISO correction integrated with the autoPROC pipeline (*19*) (STARANISO. Cambridge, United Kingdom: Global Phasing Ltd.).

The structure was solved by molecular replacement with PHASER from the PHENIX package (*9*). The resolution limit was determined based on the quality of the electron density map obtained and Rpim as major criteria. The model was manually built in COOT (*11*) and refined with Buster (*20*) (BUSTER version 2.10.4. Cambridge, United Kingdom: Global Phasing Ltd.) and PHENIX (version 1.21.1-5286). The last refinement steps were performed by refining with translation libration screw (TLS), and the model was validated by the MolProbity server (*21*). The model was refined with hydrogens in the riding position. Hydrogens were omitted in the final deposited models. Data collection and refinement statistics for the deposited models are listed in table S8.

System preparation for QM / MM calculations

We performed quantum mechanics/molecular mechanics (QM/MM) calculations using the Orca 5.0.4 quantum chemistry package (*22*) to characterize chemical properties of the experimental structures. The CODH and ACS subunits, corresponding to chains C and D in the full experimental structure, were isolated and independently prepared for calculations of the C- and A-clusters, respectively. CHARMM-GUI (*23-25*) was used to solvate the respective protein structures and prepare topology files with CHARMM36m (*26*) parameters for the protein and TIP3P model for water. The solvated structures were first energy-minimized at the MM level, fixing the positions of the atoms in the QM zone (defined below).

The QM zones contained all transition metal atoms and side chains of the coordinating amino acid residues. Side chain bonds were cropped at the αC-βC interface and filled with H link atoms. The QM zone of the A cluster contained also the backbone of the $Ni_d$-coordinating residues. The interaction between QM and MM zones was modeled with electrostatic embedding. To avoid over-polarization at the QM/MM boundaries, the automatic charge shift scheme implemented in Orca was used. The QM active region consisted of a layer of residues within 6 Å of the metallic clusters. Figure S7 shows the atomic models used for the QM calculation and for the definition of the active zone.

Broken-Symmetry Density Functional Theory (DFT) calculations

Atoms in the QM zone were modeled using the meta-GGA functional TPSS with a def2-TZVP basis set on every heavy atom (*27, 28*). Dispersion corrections were applied using the Grimme D3 method with Becke-Johnson damping (*29*). Exchange and correlation functionals at the meta-GGA level corrected for dispersion are expected to perform well for transition metal complex geometries (*30, 31*). As a first approximation to the magnetic interactions between metallic centers, we used the broken-symmetry formalism. This method has been successfully applied for the calculation of exchange coupling constants of transition metal polynuclear complexes, including metallic clusters in biological systems. Details on the broken-symmetry theory can be found elsewhere (*32*). Briefly, a self-consistent field calculation of the fully ferromagnetic state yields the electron density of the high-spin configuration. Then, molecular orbitals localized on selected metal centers are "flipped" to generate antiferromagnetic states. The lowest energy alignment is then selected as a first approximation to the ground state of the cluster. Finally, geometry optimizations are calculated with the selected broken symmetry configuration.

Chemical description with Quantum Theory of Atoms in Molecules (QTAIM)

All electronic states of the C- and A-clusters modeled in this work were used for a topological analysis of the electron density, following the QTAIM methodology (*33*). Molecular orbitals obtained by Orca were converted to Molden files and analyzed in Multiwfn (*34*). All critical points were calculated, characterized and classified. Bond paths were also generated to obtain the molecular graph of the clusters.

QM/MM data analysis

C-cluster: the formal oxidation states of the metals in the two major catalytic intermediates $C_{red1}$ and $C_{red2}$ of the C cluster are controversial, particularly for the $C_{red2}$ species. Traditionally, $C_{red1}$ is represented as $[3Fe\text{-}4S]^{-}$-$Ni^{2+}$-$Fe^{2+}$-$OH_2$, whereas the 2-electron reduced $C_{red2}$ intermediate has been identified with $[3Fe\text{-}4S]^{-}$-$Ni^{"0"}$-$Fe^{2+}$-$OH_2$ (*35*). Without density in the CryoEM maps indicating additional ligands to the metallic cluster, the geometry optimizations were calculated in the apo state, using the same formal oxidation states as the corresponding $C_{red1}$ and $C_{red2}$ intermediates, here labeled as $C_{red1(-OH)}$ and $C_{red2(-OH)}$. We performed geometry optimizations using both oxidation states and we also included a third, fully oxidized cubane $[3Fe\text{-}4S]^{+1}$-$Ni^{+2}$-$Fe^{2+}$ for comparison. These three electronic states are separated by two electrons each. The optimized structures are shown in figure S8. For the generation of the broken symmetry states, we followed a similar approach to the standard, isoelectronic $[4Fe\text{-}4S]^{+}$ cluster. First, we performed a self-consistent field calculation with a fully ferromagnetic (high spin) alignment. Then, local spins were flipped to generate antiferromagnetic alignments, with a total magnetic moment $M_s = \frac{1}{2}$, consistent with the experimental ground state. Finally, geometries were optimized for the lowest-energy broken symmetry state. Table S3 presents the energies of the antiferromagnetic alignments with respect to the high spin configuration.

A-cluster: relevant reaction intermediates in the acetyl-CoA synthesis catalyzed by the A-cluster have been characterized recently with spectroscopic methods (*36*). According to this description, within the reaction cycle, the $A_{red}$ state has an available coordination site waiting to bind either carbon monoxide or a methyl group from the CoFeSP protein. Since there is no density in the cryo-EM map suggesting an additional ligand coordinated to $Ni_p$, geometry optimizations were calculated in this low-valence state for $Ni_p$. The accepted formal oxidation states of $A_{red}$ consist of an EPR silent $[4Fe\text{-}4S]^{2+}$ cluster connected to a paramagnetic $Ni^{+}$ species, yielding a total spin of $S=\frac{1}{2}$, as demonstrated by EPR measurements (*36*). Therefore, we explored only the broken symmetry states consistent with a total $M_s = \frac{1}{2}$. The usual description of the $[4Fe\text{-}4S]^{2+}$ cluster, within the broken symmetry framework, consists of two $[2Fe\text{-}2S]^{+}$ dimers, where every iron ion within the dimer is ferromagnetically aligned to yield a mixed valence pair $Fe^{+2.5}$-$Fe^{+2.5}$ and a total spin of S=9/2. Then, the two dimers antiferromagnetically align to generate the final total spin S=0. Therefore, we selected all the possible pairs of iron dimers in the [4Fe-4S] cluster and performed local spin flips to generate the relevant broken symmetry states. Table S7 shows the results of the energy of every broken symmetry state with respect to the high spin alignment. Geometry optimization was performed in the lowest-energy broken symmetry configuration.

Sequence conservation analysis

The sequences of the ACS (WP_023162339.1), CoFeSP large (WP_023162340.1) and small subunit (WP_013240362.1) were used as query for a BLAST P analysis in the RefSeq database (*37*). This database was used to limit the redundancy of sequences from similar species. Only results exhibiting at least 80% coverage were selected, yielding 361, 417 and 408 sequences for the three query sequences, respectively, covering bacterial and archaeal enzymes. Sequences with more than 80% identity were removed using CD-HIT (*38*), yielding 103, 188 and 190 sequences, respectively. The sequences were aligned by Clustal Omega (*39*) and the residue conservation images were constructed for structurally important residues of *Ca*CODH/ACS using Weblogo 3 (*40*) (version 3.7.12) based on the multiple sequences alignment.

CoFeSP large subunit, *C. autoethanogenum*

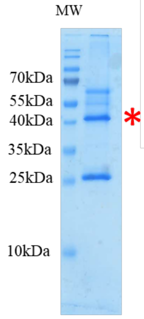


MALTGLNIFK LTPKKNCKDC GFPTCLAFSM KVAAGAVEIG KCPHMSDEAM EKLAEATAPI MKTITIGKGD
NEYKLGGETV LFRHEKTFVN RNRFAVAFSD SMDDAEVDAK IQHIKDVDYV RIGEQMKTEF AAIKYAGNKD
KYLALINKIK ASGVKVAYAL VCEDVAVMKE ALPLVKDENP LVYGANKDNF KEMVELVKGD KLALGVKADG
LEALYGLVEE IQKLGYKNLV LDPGGKSIKE AFENTVQIRR INIEGQDRTF GYPSIIFLDE LTKADKFMEV
ALSTLFTLKY GSLLVLSDMD YSRALPLYSI RQNVFTDPQK PMTVDLGIHG INNPDENSPV LCTVDFALTY
FLVSGEVERS KVPVWMVIPD AGGYSVLTSW AAGKFTGAAI ADEIKKCGIA EKTKNRTLLI PGKVAVLKGE
LEELLPDWNI VISSTEAMFI PKLLKELTAK

| Protein name | Gene name | Acc No | MW | Mass Spectrometry Analysis matched | | |
|---|---|---|---|---|---|---|
| | | | | N of unique peptides | N of spectra | Sequence coverage |
| CoFeSP large subunit | *acsC* | WP_023162340.1 | 50kDa | 32 | 951 | 66% |

**Fig. S1. Mass spectrometric identification of CoFeSP large subunit purified from *C. autoethanogenum*.** The large subunit was used to identify the complex as it should always be found in complex with the small subunit. Left: Coomassie-stained SDS gel lane containing purified CoFeSP. The gel band analyzed by MS is designated with a red asterisk, and protein molecular weight markers are shown on the left. Top: distribution of peptides detected by MS for the CoFeSP large subunit in the analyzed gel band. MS-matched peptides are highlighted in yellow, and amino acids detected with modifications (oxidation, acetylation) are marked in green. Bottom: MS-identification details for CoFeSP large subunit in the corresponding band.

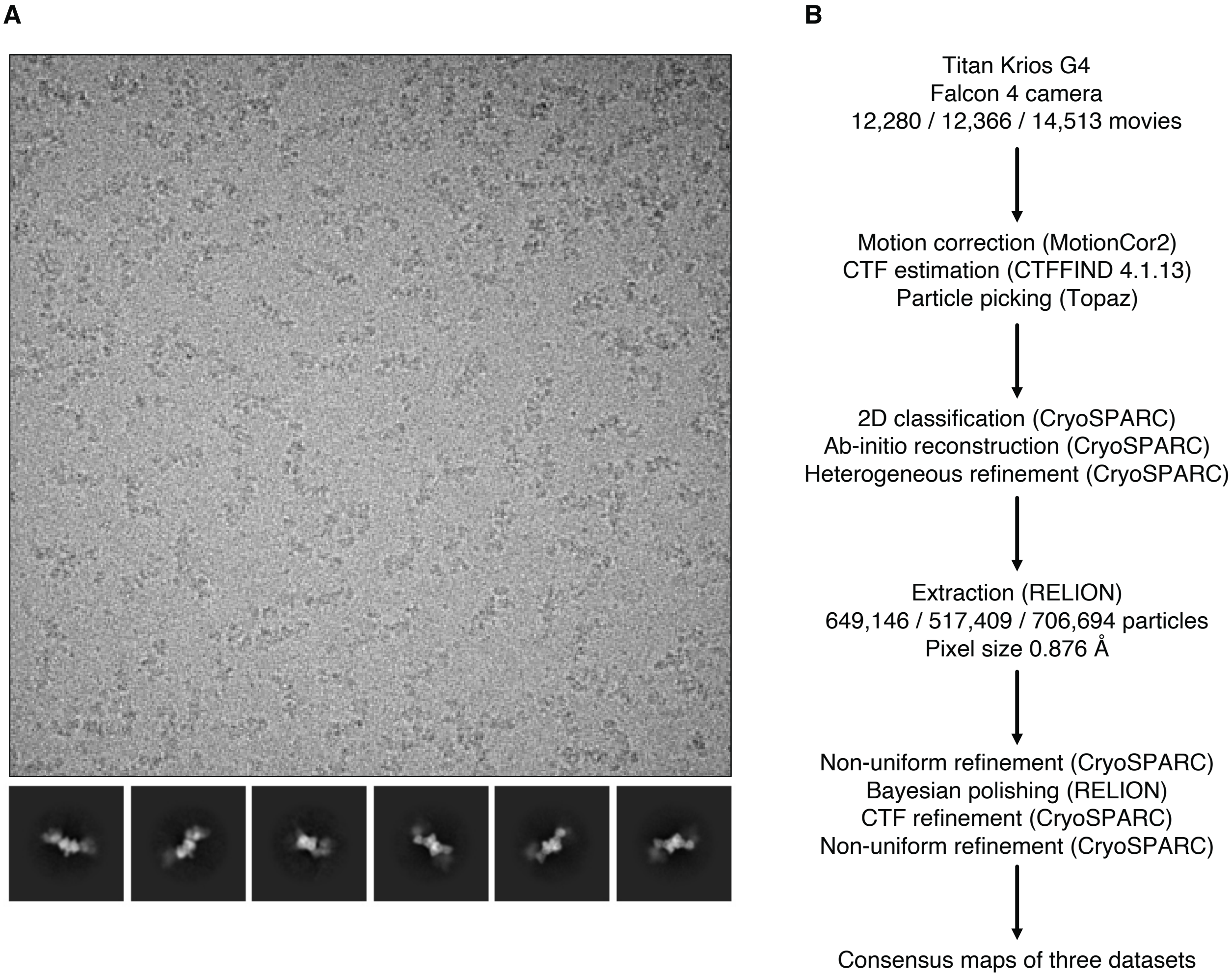


**Fig. S2. Cryo-EM processing workflow until consensus refinements. (A)** A representative micrograph and representative 2D classes. The micrograph was lowpass-filtered to 20 Å and displayed with 0 sigma contrast. **(B)** Single-particle cryo-EM processing workflow until completion of consensus refinements.

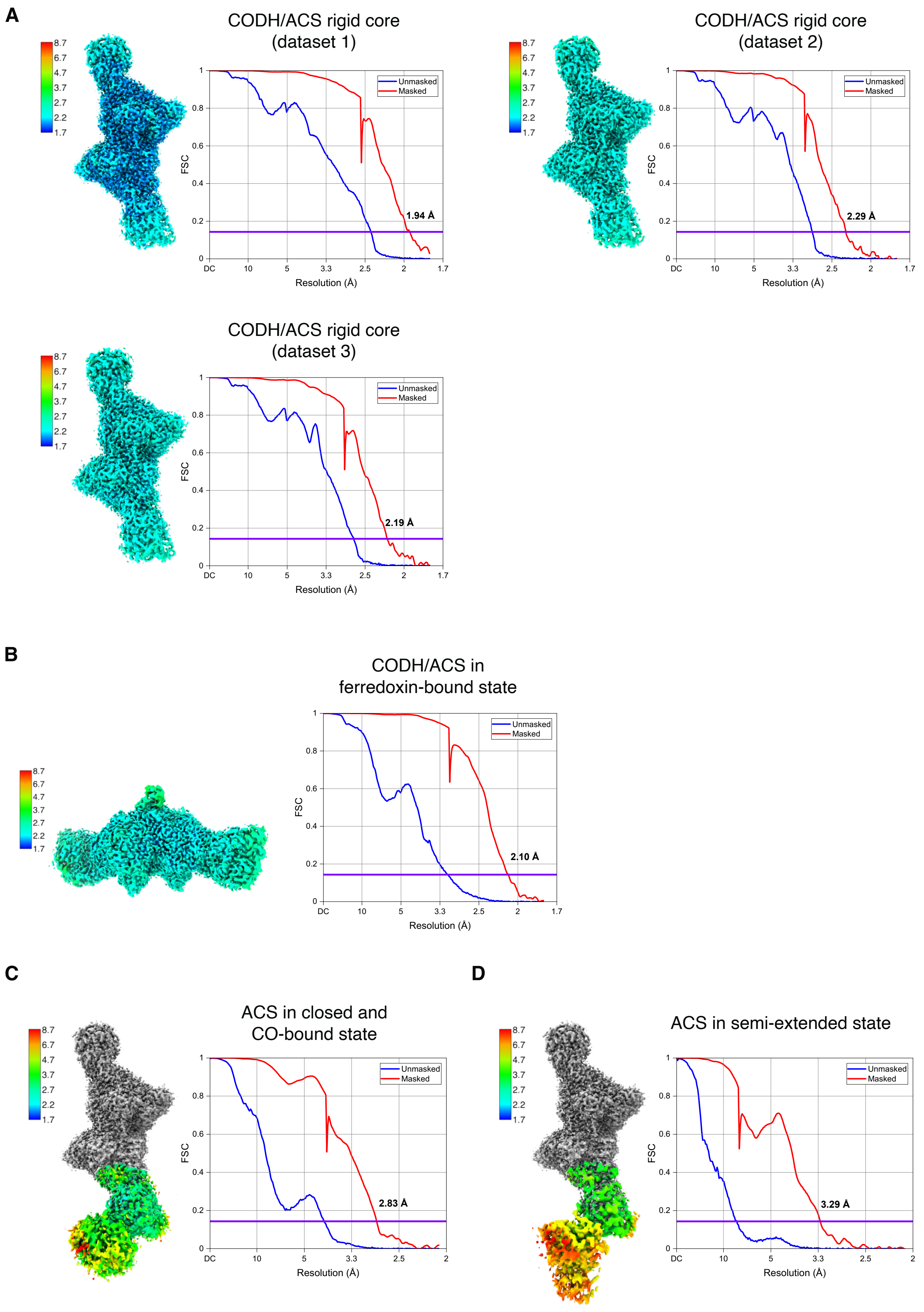
A
CODH/ACS rigid core
(dataset 1)
Unmasked
Masked
1.94 Å
FSC
Resolution (Å)
CODH/ACS rigid core
(dataset 2)
Unmasked
Masked
2.29 Å
FSC
Resolution (Å)
CODH/ACS rigid core
(dataset 3)
Unmasked
Masked
2.19 Å
FSC
Resolution (Å)
B
CODH/ACS in
ferredoxin-bound state
Unmasked
Masked
2.10 Å
FSC
Resolution (Å)
C
ACS in closed and
CO-bound state
Unmasked
Masked
2.83 Å
FSC
Resolution (Å)
D
ACS in semi-extended state
Unmasked
Masked
3.29 Å
FSC
Resolution (Å)

**Fig. S3. Local resolution estimates and Fourier shell correlation (FSC) curves.** Local resolution estimation was performed in cryoSPARC and visualized in ChimeraX. FSC between independent half-maps of the CODH/ACS rigid cores **(A)**, the CODH/ACS in ferredoxin-bound state **(B),** closed state **(C)** and semi-extended state **(D)** are plotted. The reported resolutions were determined using gold-standard FSC at 0.143 threshold, indicated by purple horizontal lines, following refinements in cryoSPARC. The best-resolved CODH/ACS rigid core (dataset 1) was used for further data analysis.

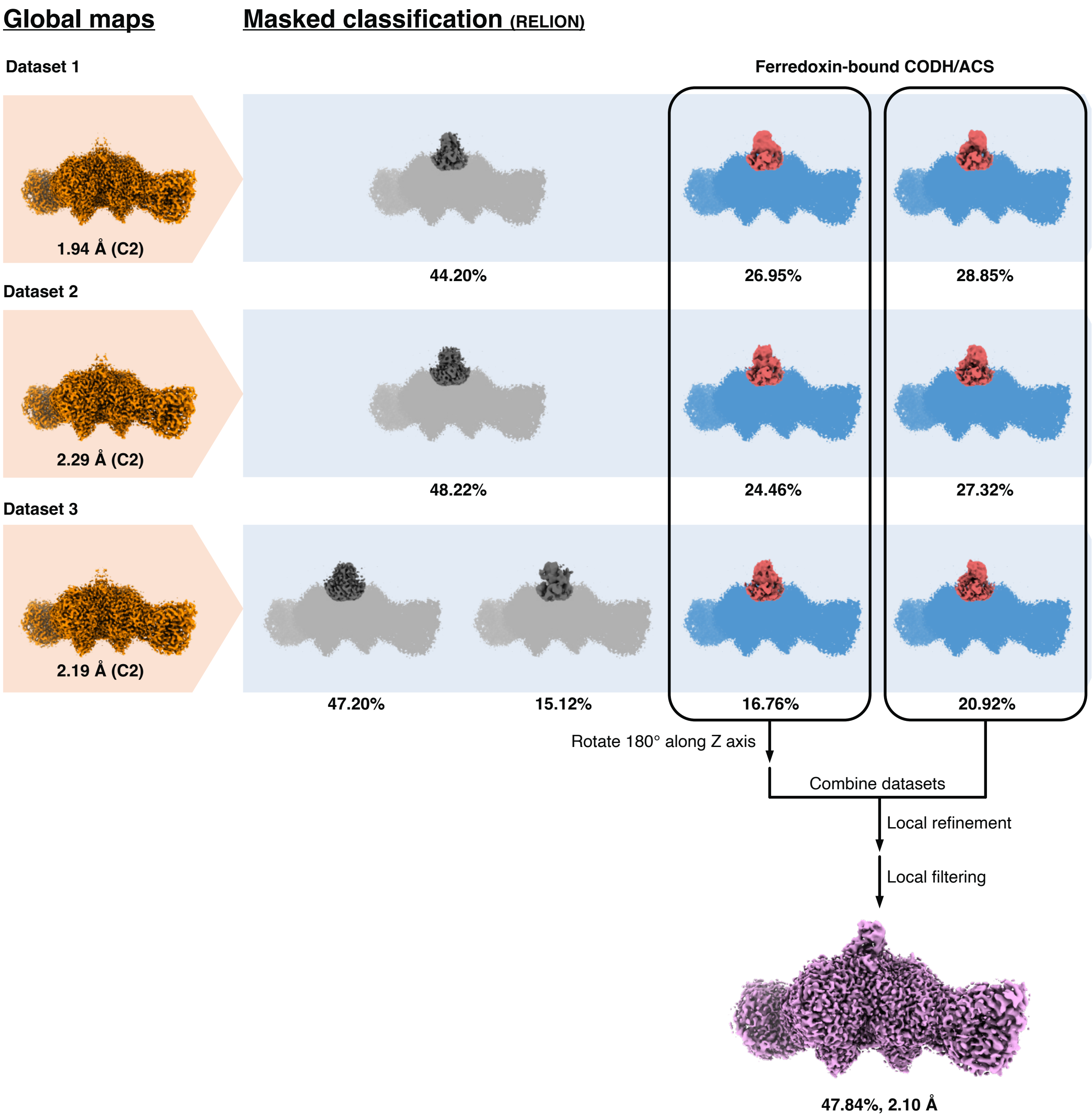


**Fig. S4. Cryo-EM processing workflow for CODH/ACS in ferredoxin-bound state.** The reported resolutions were determined using gold-standard FSC at 0.143 threshold after the refinements in cryoSPARC. The proportions of particles contributing to each 3D class and the final map are also provided.

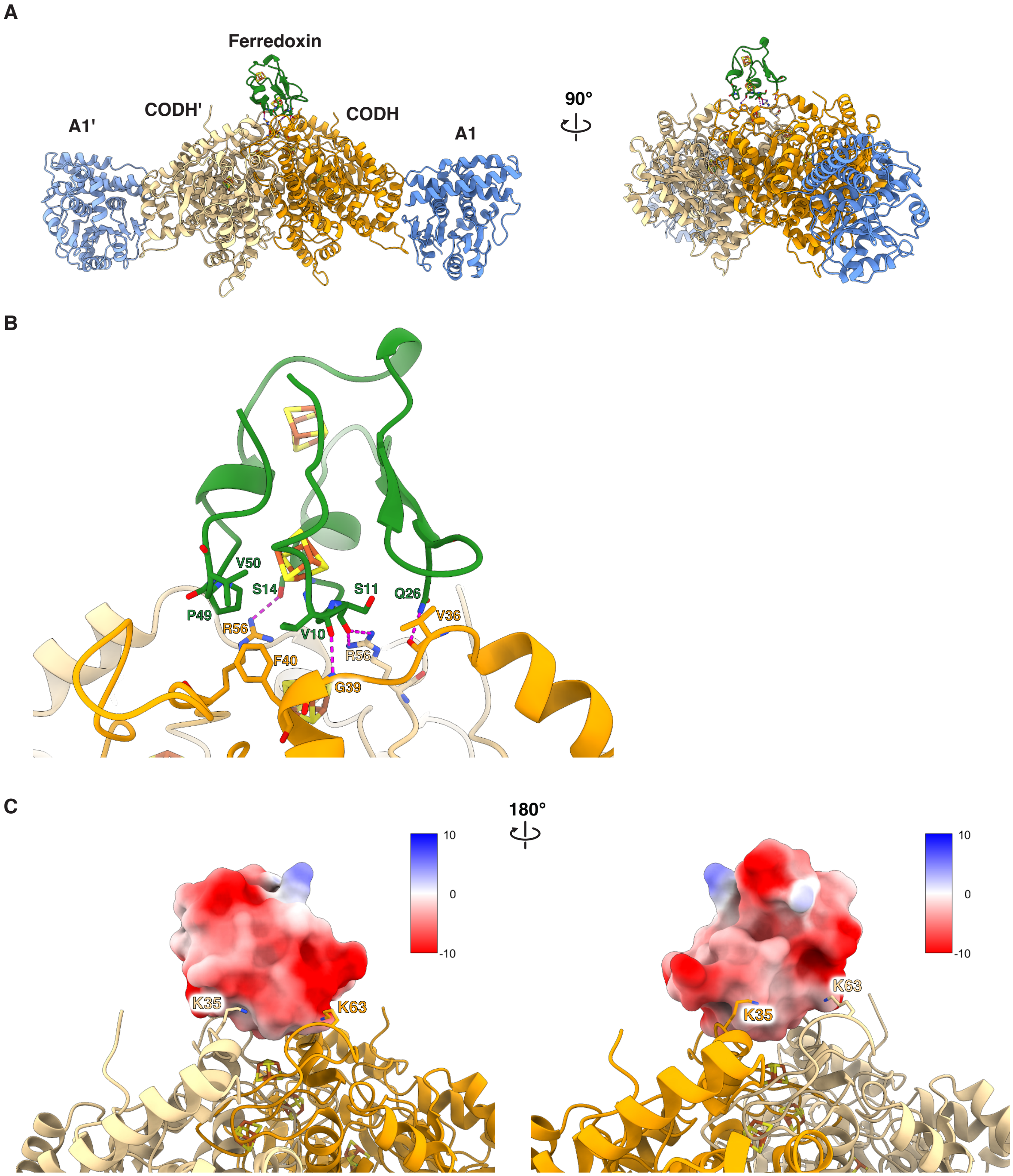


**Fig. S5. Interaction between CODH and ferredoxin. (A)** Overviews of the CODH/ACS rigid core complexed with ferredoxin (forest green), using the same color scheme as in Fig. 1 for CODH/ACS. **(B)** A close-up view of the interface, highlighting the hydrogen bonding (magenta dashed lines) and hydrophobic interactions. **(C)** Electrostatic interactions likely further stabilize ferredoxin, shown as a molecular surface colored by the electrostatic potential (in units of kcal/(mol•*e*) at 298 K) in ChimeraX (*16*). CODH residues involved in these interactions are highlighted.

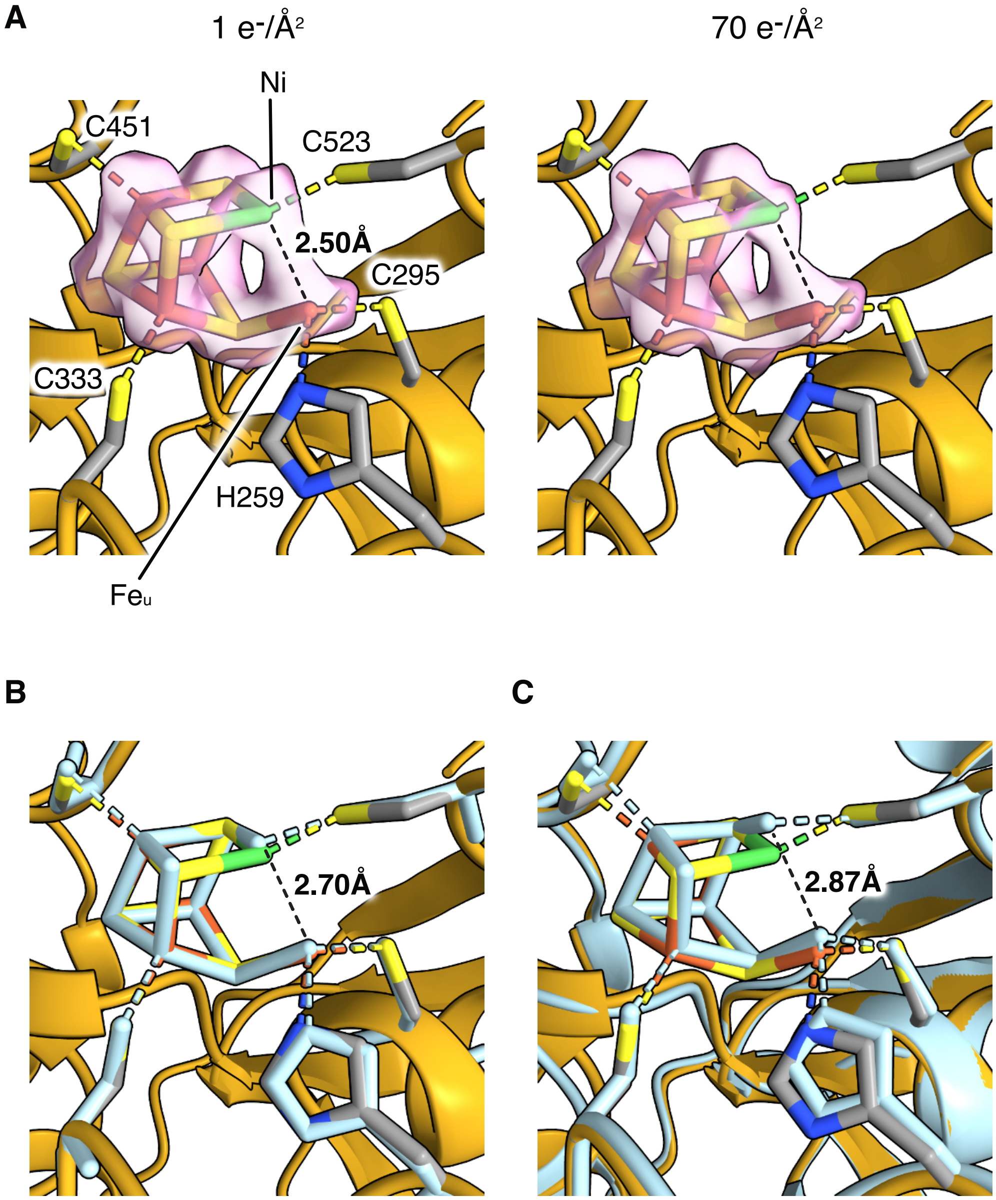


**Fig. S6. Close-up views of the C-cluster. (A)** The cryo-EM map densities of the C-cluster, reconstructed for different total electron doses, do not reveal additional density due to bound ligands or any notable difference in the Ni-$Fe_u$ distance. The map corresponding to an electron fluence of 1 $e^-/Å^2$, equivalent to a dose of 3.7MGy (*41*), was generated via early frame reconstruction (see Methods). The CODH (class 3Cβ, chain B) is shown as gold ribbon, while the C-cluster and ligating residues are depicted as sticks colored according to their elemental types. Cryo-EM densities of the C-cluster are shown as light pink surfaces, displayed at 10.9 sigma (left panel) and 20.1 sigma (right panel), respectively. The observed distance between Ni and the $Fe_u$ (black dashed lines) is relatively short in the current study (set at 2.5 Å), when compared with other experimental structures of the C-cluster: the as-isolated *Ca*CODH dimer (*1*) (**B**, PDB 6YU9, chain B; light blue; 1.90 Å resolution) and the CO-treated *Mt*CODH/ACS (*42*) (**C**, PDB 6X5K, chain A; light blue; 2.47 Å resolution). Residues 479-492 (*Ca*CODH numbering) or residues 498-511 (*Mt*CODH numbering) are omitted for clarity.

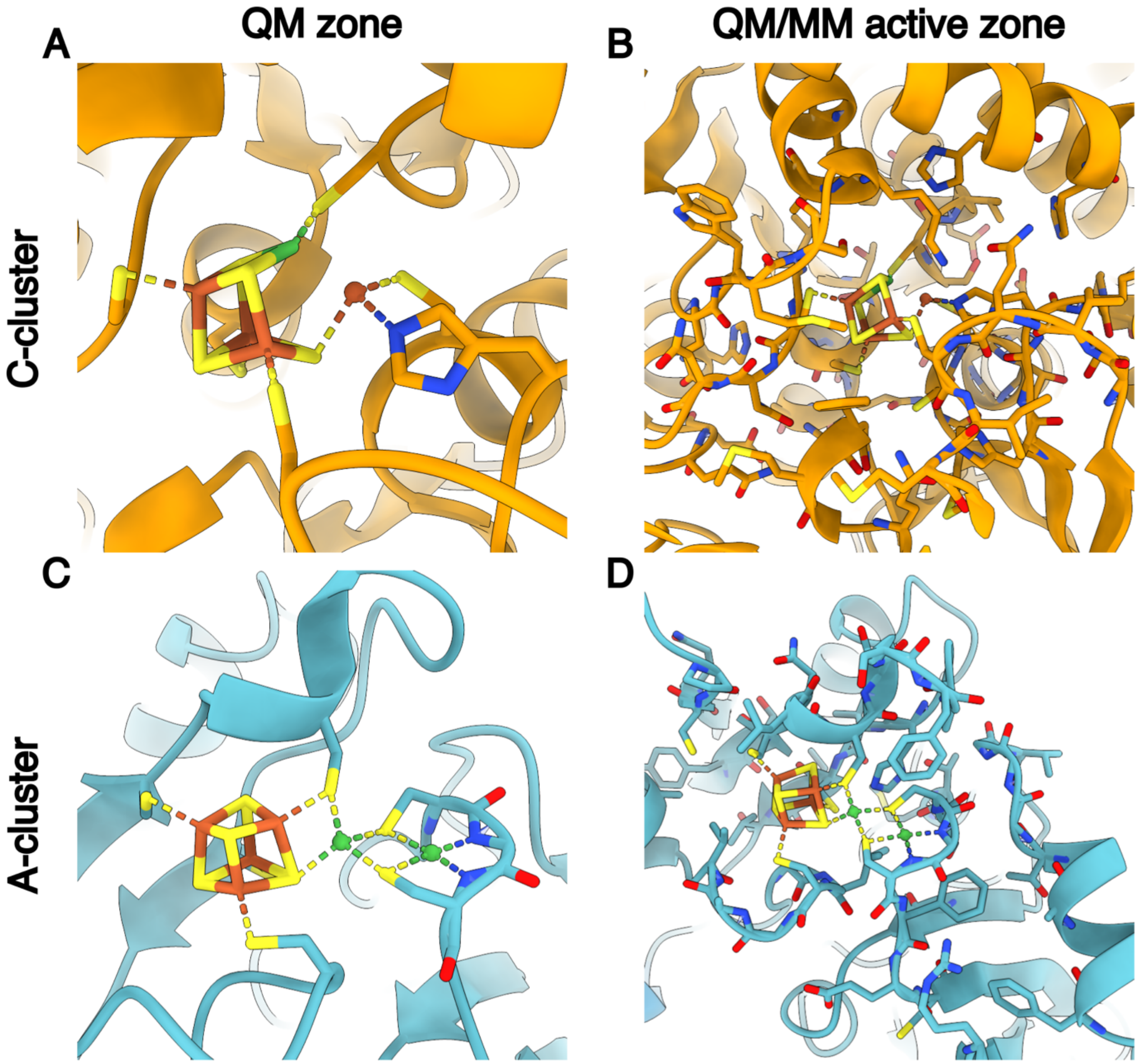


**Fig. S7. Definition of the QM and MM active zones.** Experimental structures showing the atoms contained in the QM zone and in the active QM/MM region, for the C-cluster **(A, B)** and for the A-cluster **(C, D)**. All non-hydrogen atoms contained in each zone are shown with stick models and the rest of the protein is displayed as ribbons. Each protein chain was used for geometry optimization separately.

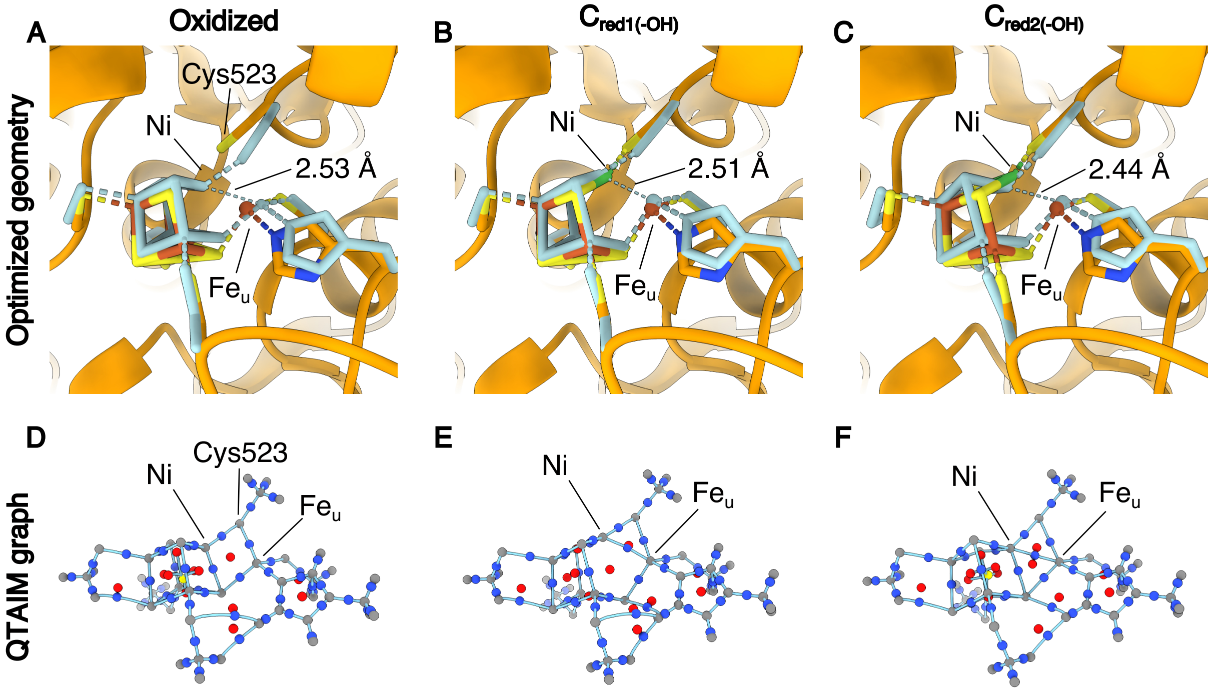


**Fig. S8. Geometry-optimized structure and QTAIM analysis of the C-cluster.** Predicted structures of the C-cluster after optimization at the TPSS-D3(BJ)/def2-TZVP level of theory for three different formal oxidation states: oxidized, $[3Fe\text{-}4S]^{+}$-$Ni^{2+}$-$Fe^{2+}$ **(A)**, $C_{red1(-OH)}$ $[3Fe\text{-}4S]^{-}$-$Ni^{2+}$-$Fe^{2+}$ **(B),** and $C_{red2(-OH)}$ $[3Fe\text{-}4S]^{-}$-$Ni^{\text{"0"}}$-$Fe^{2+}$ **(C)**. Experimental starting structures are shown in orange, and DFT-optimized models are superimposed and colored in blue. **(D-F)** The molecular graphs are calculated from the topology of the electron density from the corresponding oxidation state. Critical points are colored according to their classification: nuclear in gray, bond in blue, ring in red, and cage in yellow. Calculated bond paths are represented as continuous light blue lines.

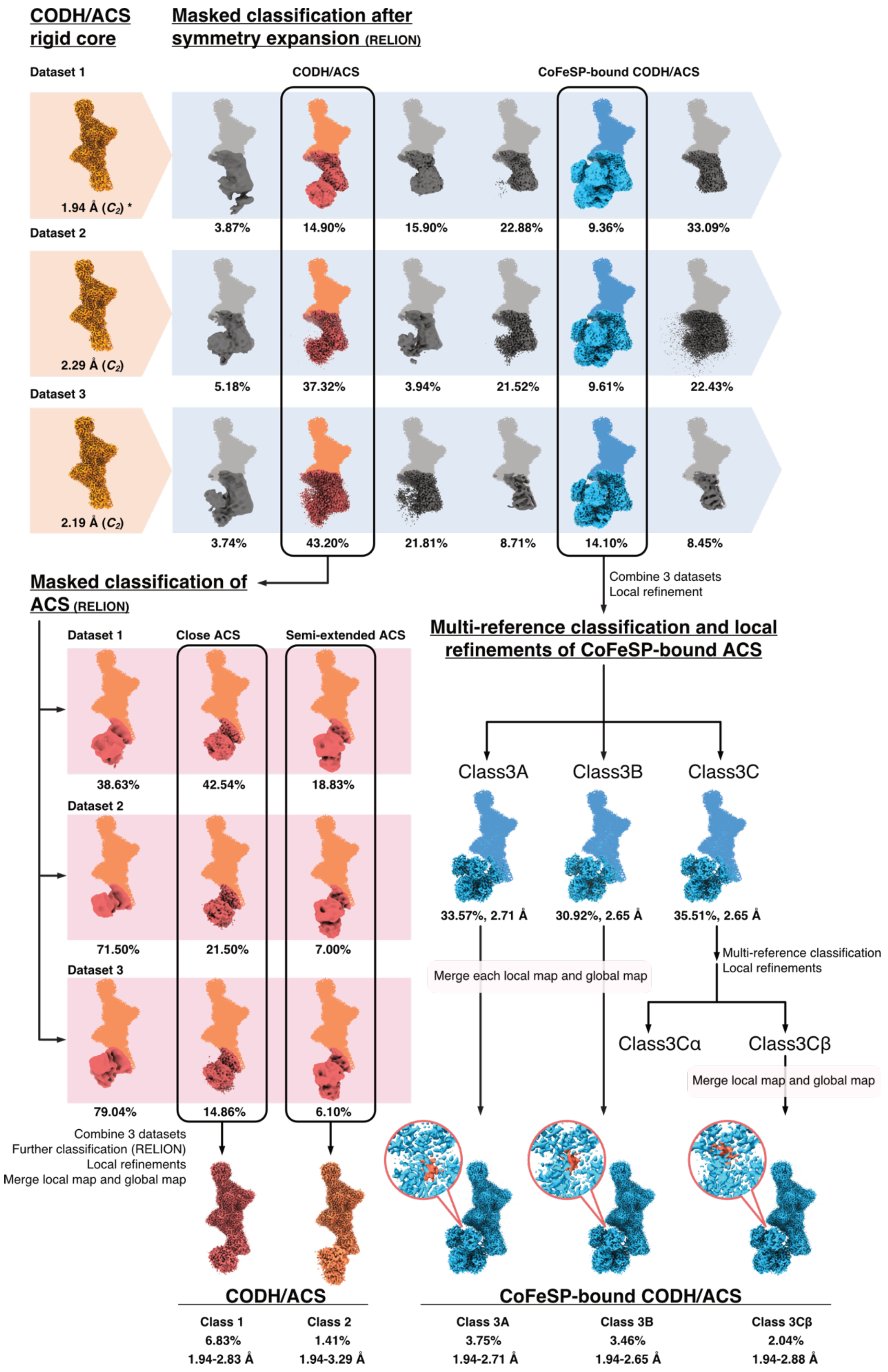
CODH/ACS
rigid core
Masked classification after
symmetry expansion (RELION)
Dataset 1
CODH/ACS
CoFeSP-bound CODH/ACS
1.94 Å (C2) *
3.87%
14.90%
15.90%
22.88%
9.36%
33.09%
Dataset 2
2.29 Å (C2)
5.18%
37.32%
3.94%
21.52%
9.61%
22.43%
Dataset 3
2.19 Å (C2)
3.74%
43.20%
21.81%
8.71%
14.10%
8.45%
Masked classification of
ACS (RELION)
Combine 3 datasets
Local refinement
Multi-reference classification and local
refinements of CoFeSP-bound ACS
Dataset 1
Close ACS
Semi-extended ACS
38.63%
42.54%
18.83%
Dataset 2
71.50%
21.50%
7.00%
Dataset 3
79.04%
14.86%
6.10%
Class3A
Class3B
Class3C
33.57%, 2.71 Å
30.92%, 2.65 Å
35.51%, 2.65 Å
Multi-reference classification
Local refinements
Merge each local map and global map
Class3Cα
Class3Cβ
Merge local map and global map
Combine 3 datasets
Further classification (RELION)
Local refinements
Merge local map and global map
CODH/ACS
Class 1
6.83%
1.94-2.83 Å
Class 2
1.41%
1.94-3.29 Å
CoFeSP-bound CODH/ACS
Class 3A
3.75%
1.94-2.71 Å
Class 3B
3.46%
1.94-2.65 Å
Class 3Cβ
2.04%
1.94-2.88 Å

**Fig. S9. Cryo-EM processing workflow for the CODH/ACS and the CoFeSP-bound CODH/ACS.** The reported resolutions were determined using gold-standard FSC at 0.143 threshold after the refinements in cryoSPARC. Resolutions of the composite maps, which span a range derived from the resolutions of the respective component maps, are also reported, alongside the proportions of particles contributing to each state. The consensus map from dataset 1, used in the preparation of composite maps, is denoted with an asterisk. We did not observe states corresponding to the 'extended' and 'hyper-extended' states described by negative-stain EM for *Mt*CODH/ACS in the absence of CoFeSP (*43*). We do not rule out the possibility that such states exist in our sample as the majority of the particles did not exhibit any clear density for the A2 and A3, which could be due to extreme flexibility.

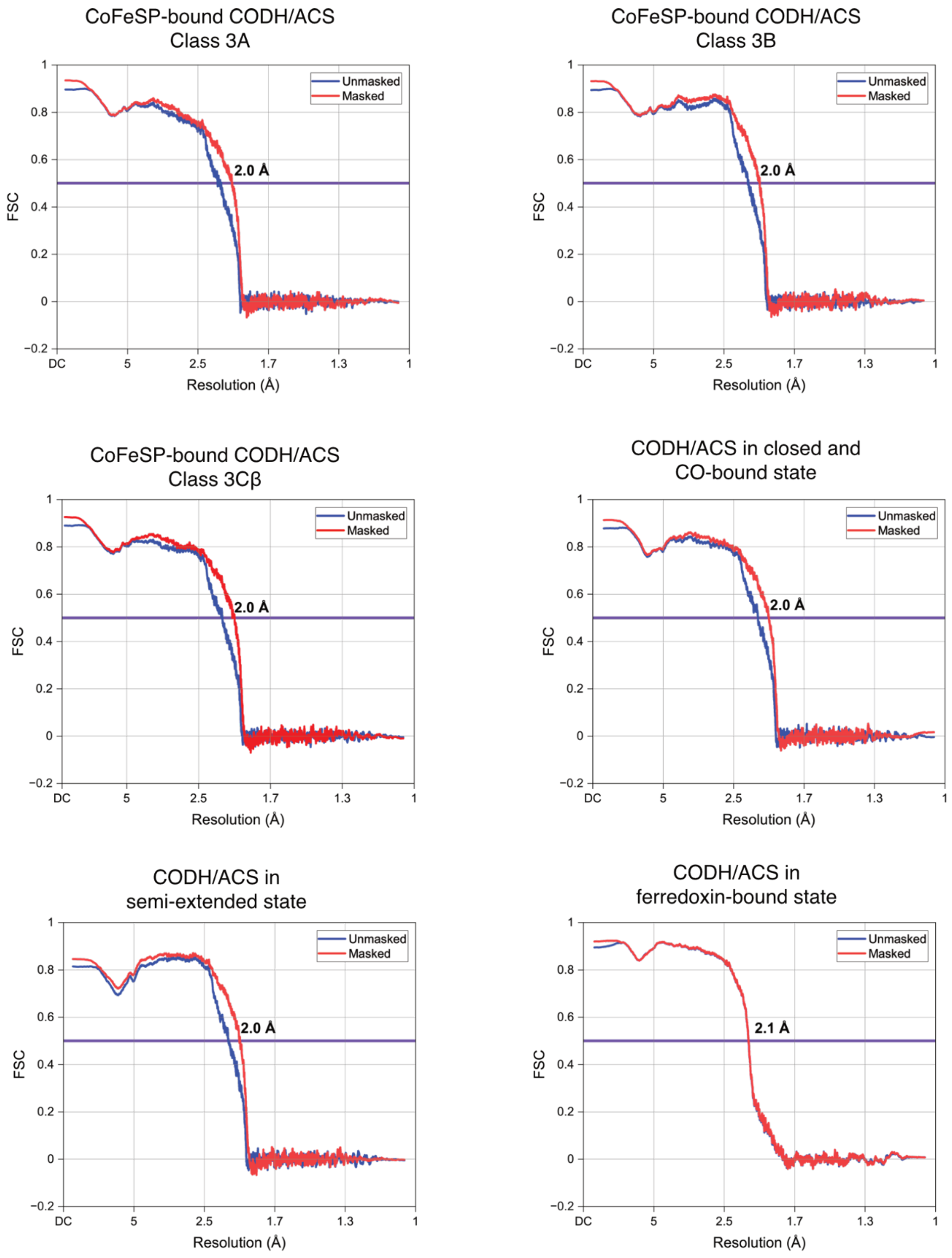


**Fig. S10. Model-map FSC curves.** FSC between refined atomic models and corresponding maps are plotted. The reported resolutions were determined using correlations at 0.5 threshold, indicated by purple horizontal lines, following validations in PHENIX.

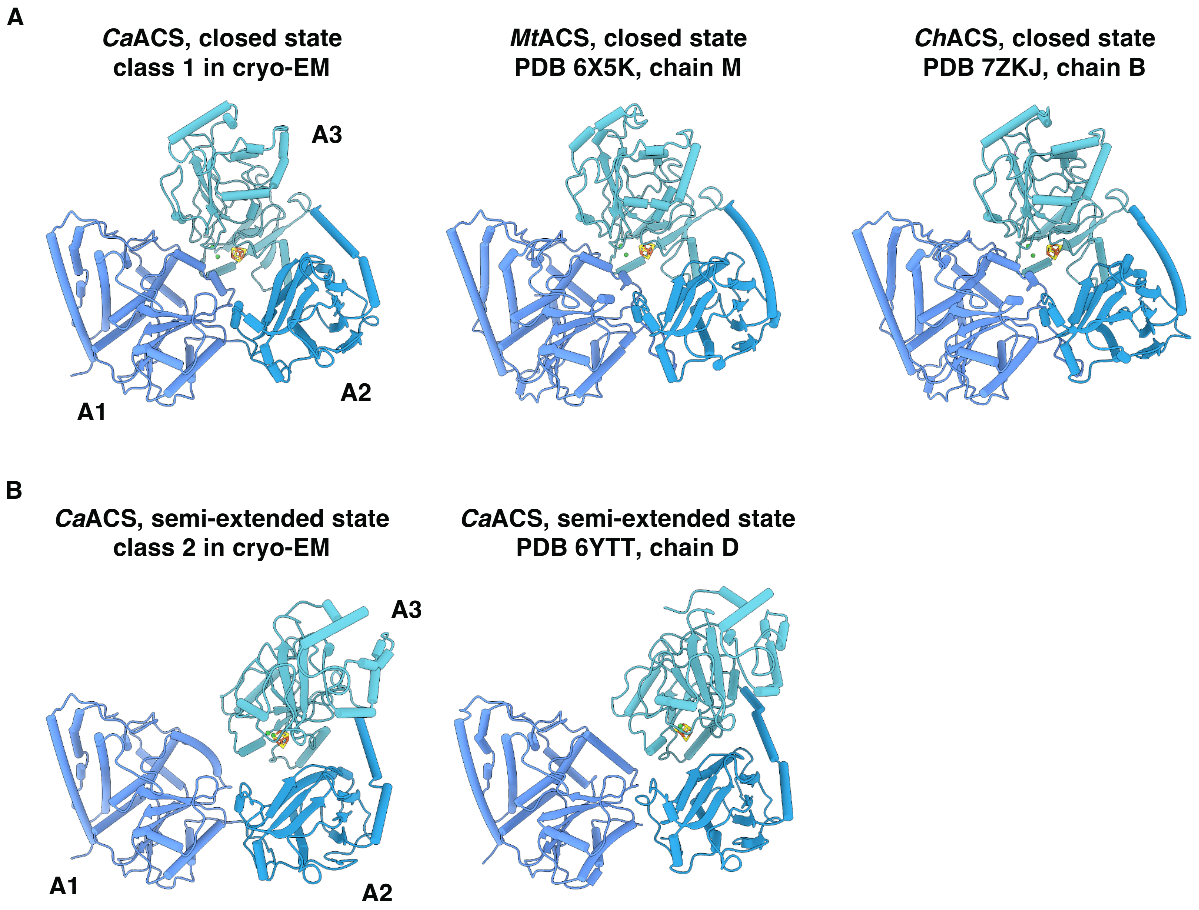


**Fig. S11. Comparison of closed and semi-extended ACS conformations from the current study with published ACS structures. (A)** Comparison of closed *Ca*ACS obtained in class 1 with closed *Mt*ACS and *Ch*ACS. **(B)** Comparison of semi-extended *Ca*ACS obtained in class 2 with a previously published semi-extended *Ca*ACS.

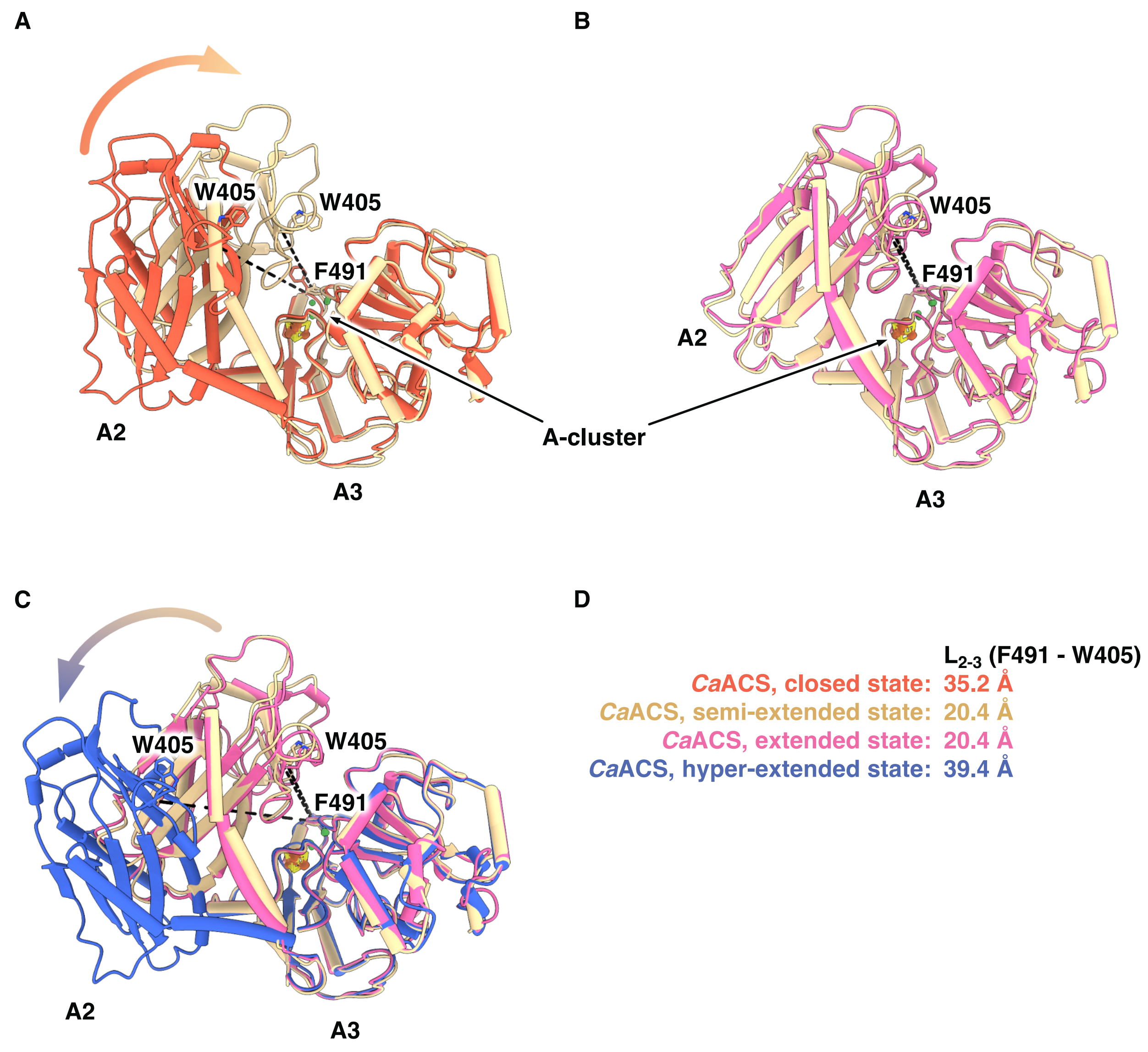


**Fig. S12. Movement between A2 and A3 across different ACS conformations. (A)** Closing of the A2-A3 interdomain space from the closed ACS to semi-extended ACS. **(B)** Semi-extended and extended ACSs share similar A2-A3 interdomain spacing. **(C)** Opening of the A2-A3 interdomain space from the semi-extended or extended ACS to hyper-extended ACS. Closed (class 1; red), semi-extended (class 2; wheat), extended (PDB 6YTT, chain A (*1*); pink) and hyper-extended ACS (class 3Cβ; blue) are aligned by the A3. For simplicity, only A2 and A3 are visualized. $L_{2-3}$ distances, measured between F491 and W405 (*Ca*ACS numbering), are indicated by dashed lines in **(A-C)**, and reported in **(D)**.

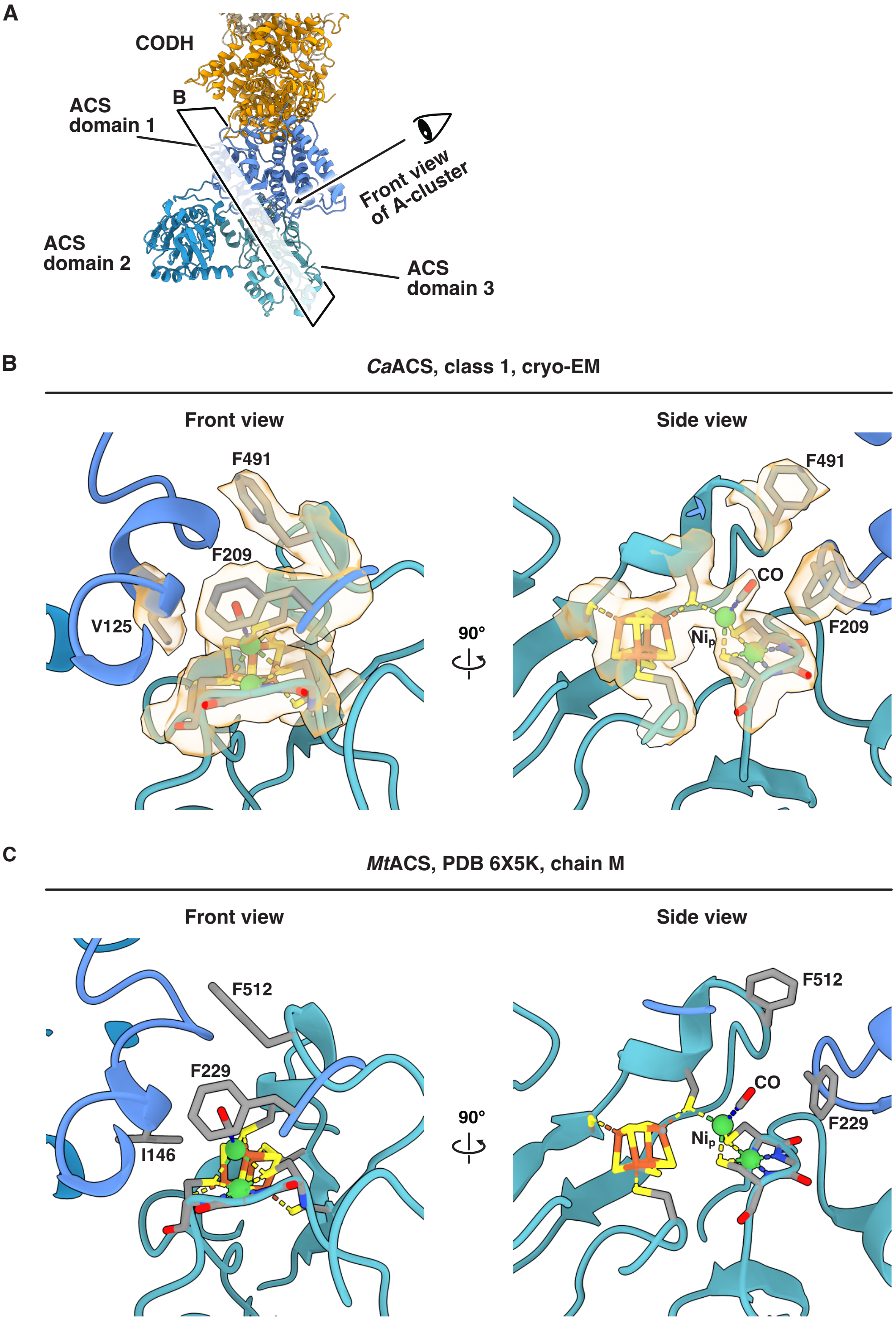


**Fig. S13. Structural similarity of ACS from different organisms in the closed state. (A)** Overview of the *Ca*CODH/ACS in the closed state. Key residues aiding in carbonylation of the $Ni_p$ exhibit similar positioning and orientation in *Ca*ACS **(B)** and *Mt*ACS (PDB 6X5K, chain M, (*42*)) **(C)**. Cryo-EM density surrounding the A-cluster of *Ca*ACS is shown as light orange surfaces, displayed at 7.93 sigma.

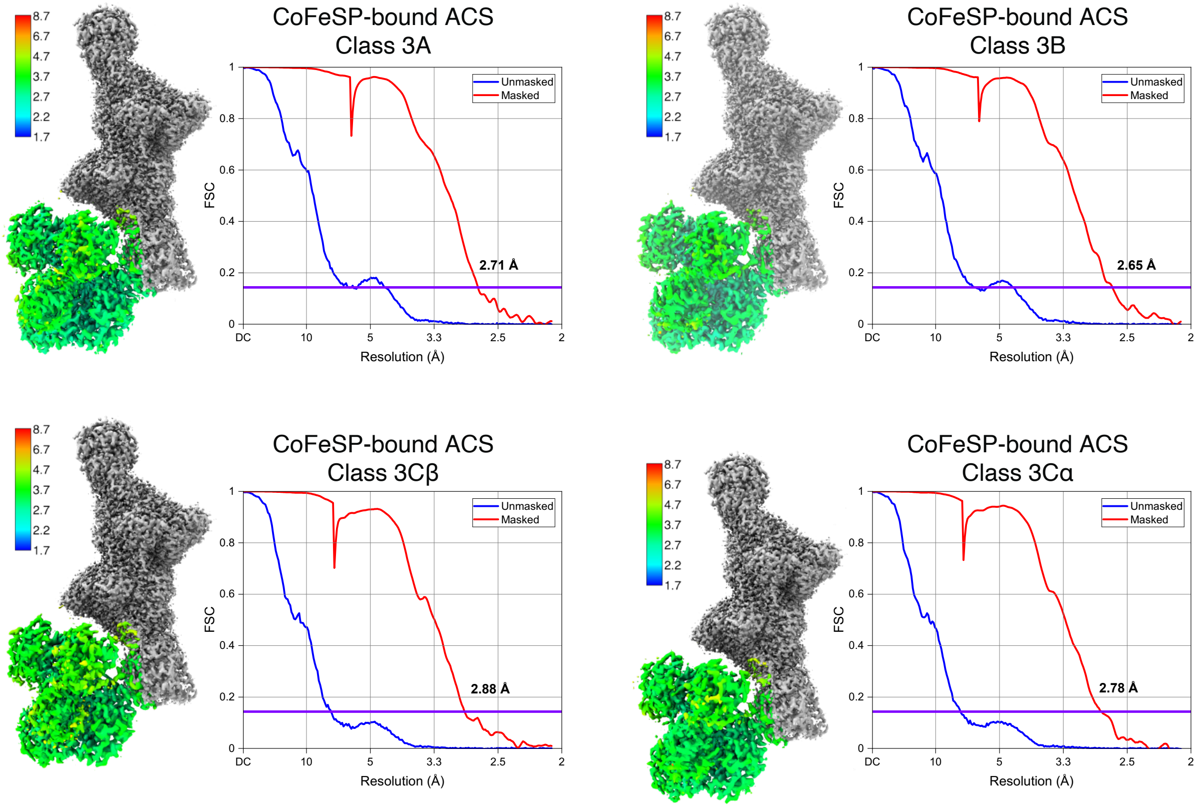


**Fig. S14. Local resolution estimates and FSC curves**. Local resolution estimation was performed in cryoSPARC and visualized in ChimeraX. FSC between independent half-maps of the CODH/ACS in various CoFeSP-bound states are plotted. The reported resolutions were determined using gold-standard FSC at 0.143 threshold, indicated by purple horizontal lines, following refinements in cryoSPARC.

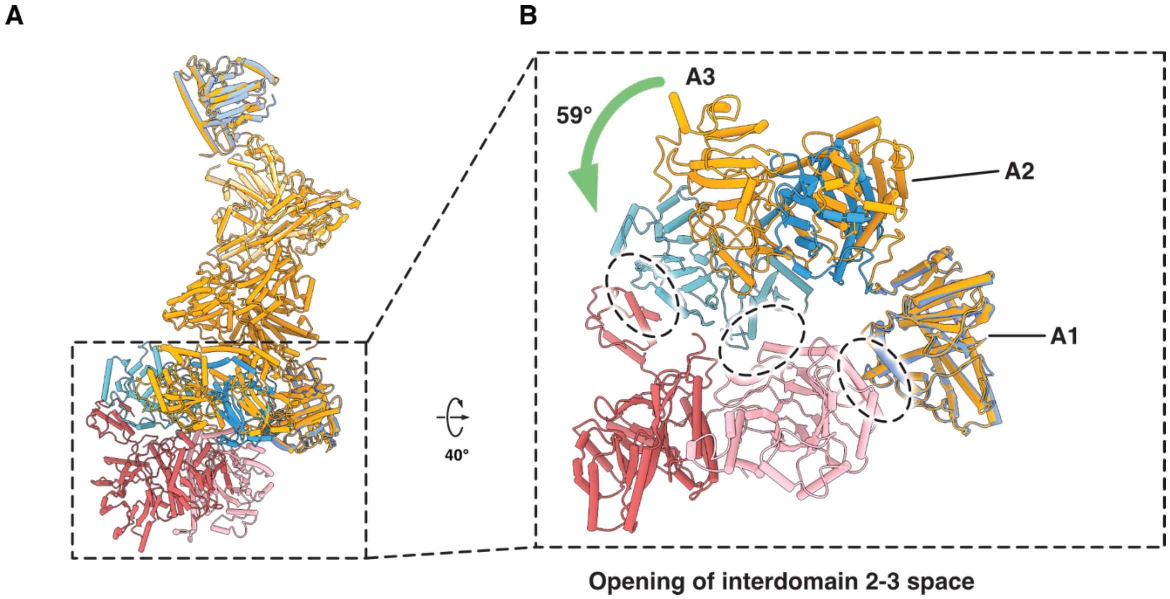


**Fig. S15. ACS must transit from the extended to the hyper-extended state in order to bind the CoFeSP. (A)** Superposition of a hyper-extended CODH/ACS complexed with CoFeSP (class 3Cβ, cryo-EM), with an extended CODH/ACS (PDB 6YTT (*1*), chain A; gold), aligned by the CODH rigid core. **(B)** A close-up view of the comparison. Interfaces between the hyper-extended ACS and CoFeSP are highlighted with dashed circles (see figs. S16-17), while motion of A3 is indicated with a green arrow. The B12 domain is omitted for clarity.

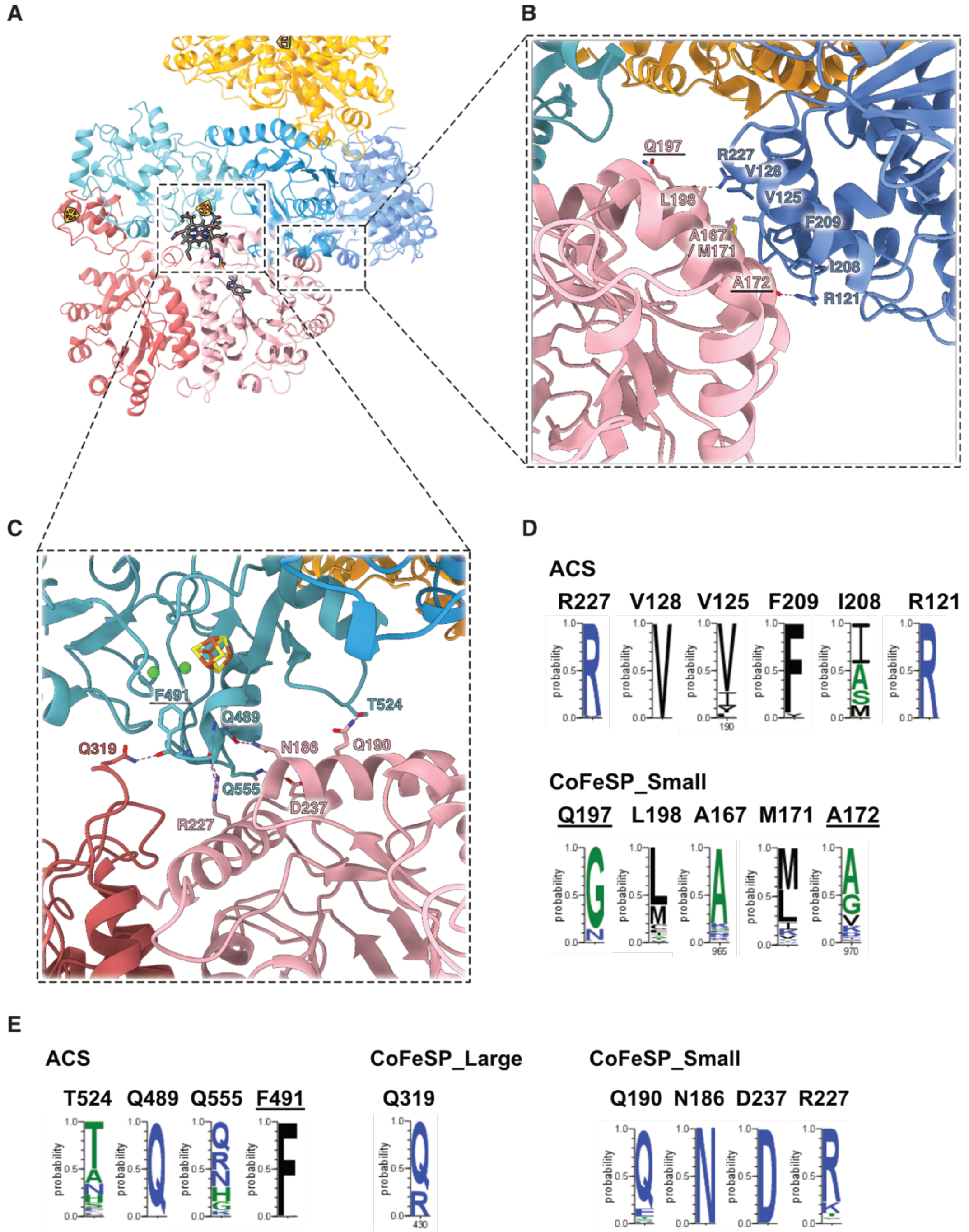


**Fig. S16 ACS:CoFeSP interaction is primarily mediated by the CoFeSP small subunit. (A)** An overview of the CODH/ACS complexed with CoFeSP. The CoFeSP establishes extensive interactions with A1 and A3. ACS:CoFeSP interaction mediated by the [4Fe-4S] cluster domain of CoFeSP is described in fig. S17. **(B)** A close-up view of the interface between A1 and CoFeSP small subunit, with the predominantly hydrophobic interactions highlighted. **(C)** Detailed view of the interface between A3 and CoFeSP, showing hydrogen bonds as magenta dashed lines. Sequence conservation analysis shows that the residues stabilizing A1:CoFeSP interaction **(D)** and A3:CoFeSP interaction **(E)** are globally conserved. Underlined residues interact via the main chain.

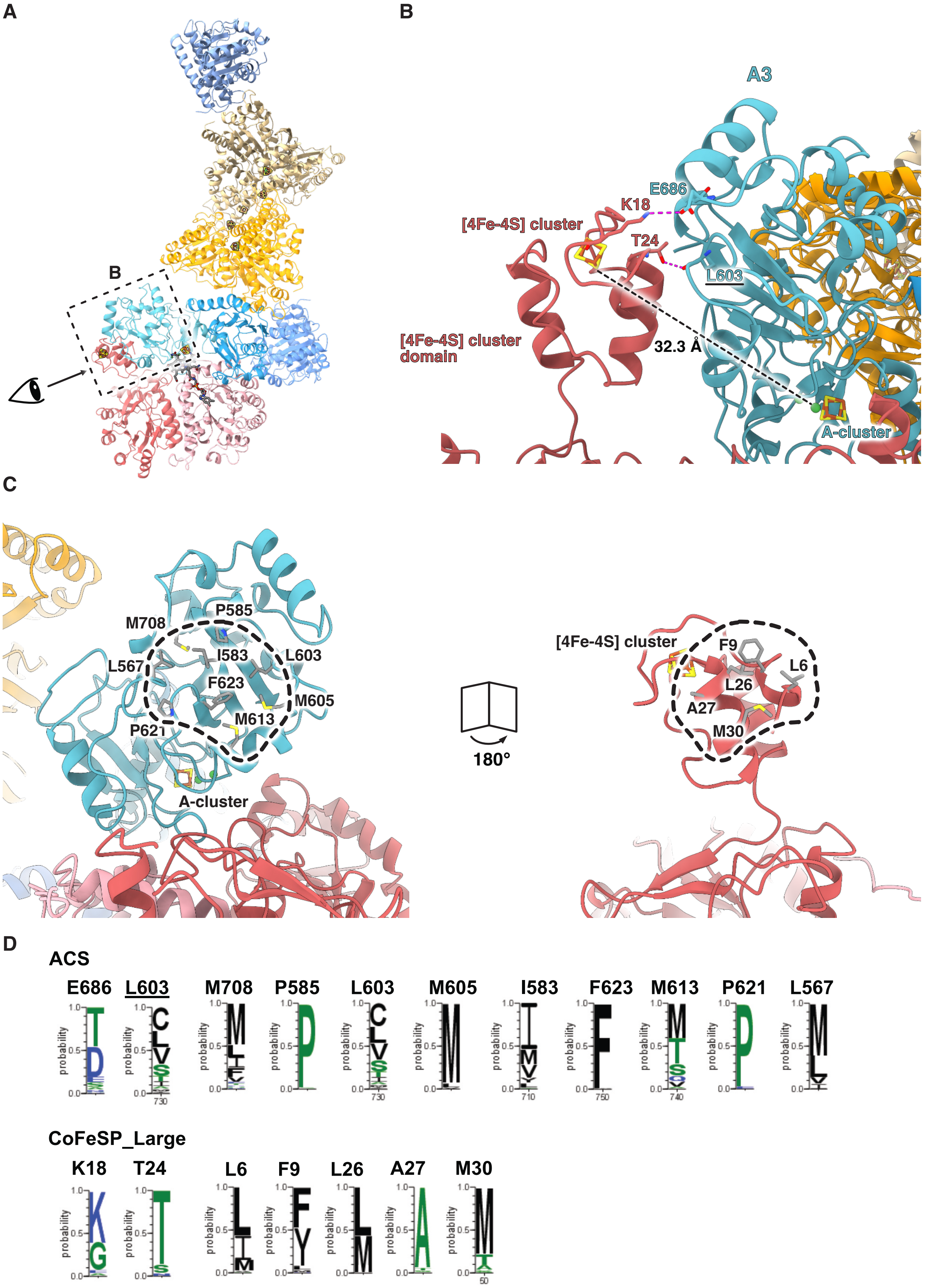
A
B
B
A3
E686
K18
[4Fe-4S] cluster
T24
L603
[4Fe-4S] cluster domain
32.3 Å
A-cluster
C
M708
P585
I583
L603
L567
F623
M605
M613
P621
A-cluster
180°
[4Fe-4S] cluster
F9
L6
L26
A27
M30
D
ACS
E686
L603
M708
P585
L603
M605
I583
F623
M613
P621
L567
CoFeSP_Large
K18
T24
L6
F9
L26
A27
M30

**Fig. S17. The [4Fe-4S] cluster domain of CoFeSP contributes to the ACS:CoFeSP interaction primarily via hydrophobic interactions. (A)** An overview of the CODH/ACS complexed with CoFeSP. **(B)** A close-up view of the interface, highlighting hydrogen bonds as magenta dashed lines, and distance measurement between the A-cluster and the [4Fe-4S] cluster as a black dashed line. **(C)** Side-by-side exposed interface, highlighting residues engaged in hydrophobic interactions. **(D)** Sequence conservation analysis shows that for most of the residues involved in the interactions, the physicochemical properties (*e.g.,* hydrophobicity) are well conserved. Underlined residues interact via the main chain.

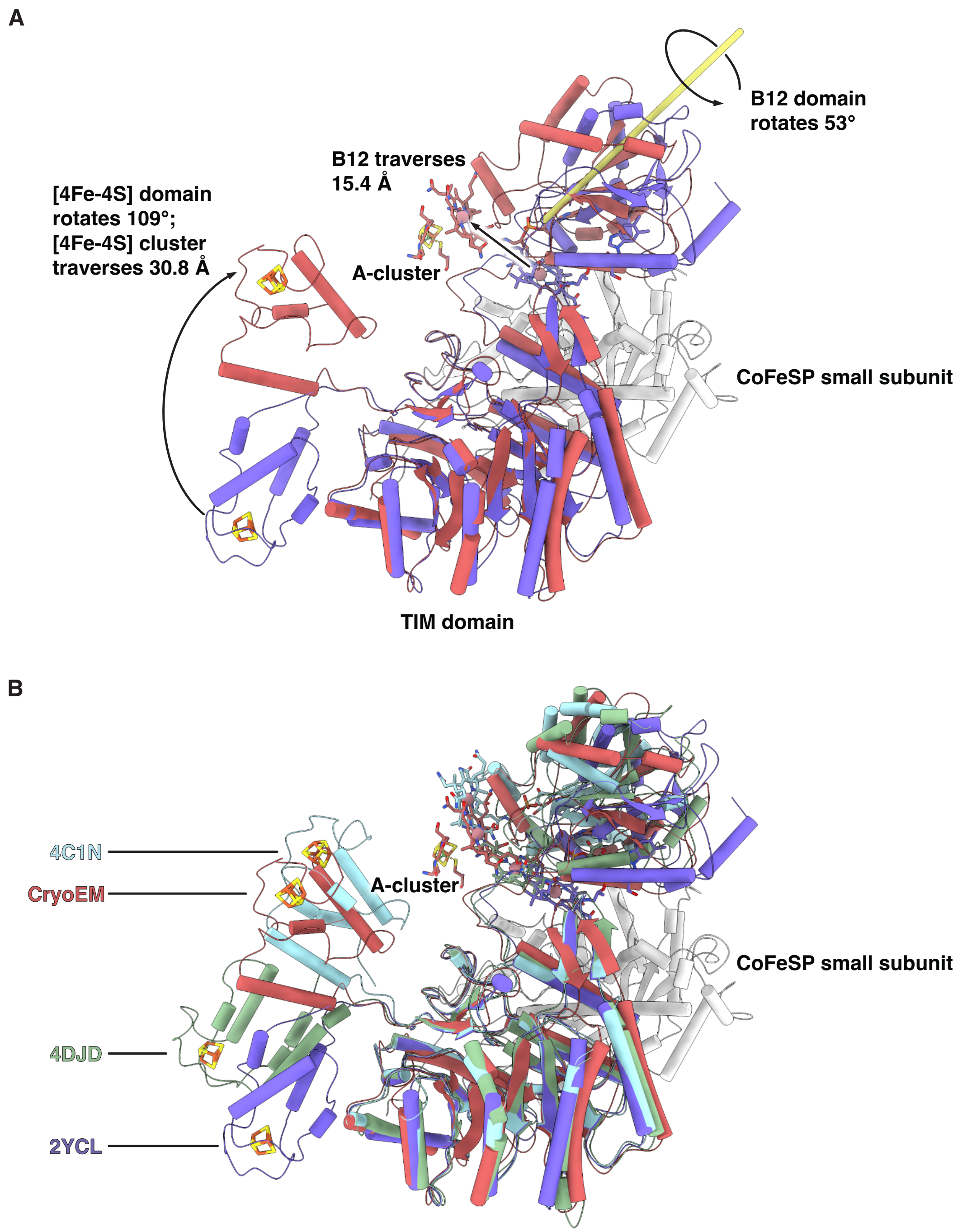


**Fig. S18. Structural plasticity of the [4Fe-4S] cluster domain and B12 domain of CoFeSP.** **(A)** Movement of these cofactor-carrying domains when bound to CODH/ACS (*Ca*CoFeSP, class 3Cβ; red), compared to a CoFeSP in resting state (*Ch*CoFeSP, PDB 2YCL (*44*); purple). **(B)** Conformational plasticity of CoFeSP, as illustrated by characterized structural states of CoFeSP across different species: a resting state (*Ch*CoFeSP, PDB 2YCL (*44*); purple), complexed with a methyl donor MeTr (*Mt*CoFeSP, PDB 4DJD (*45*); light green), a methyl acceptor ACS (class 3Cβ in our cryo-EM study, red) or a reductive activator RACo (*Ch*CoFeSP, PDB 4C1N (*46*); cyan). The CoFeSP small subunit (gray cartoon) and A-cluster (sticks, colored according to the elemental types) in the class 3Cβ are illustrated for context. Structures are aligned according to the triosephosphate isomerase (TIM) barrel domain of the CoFeSP large subunit.

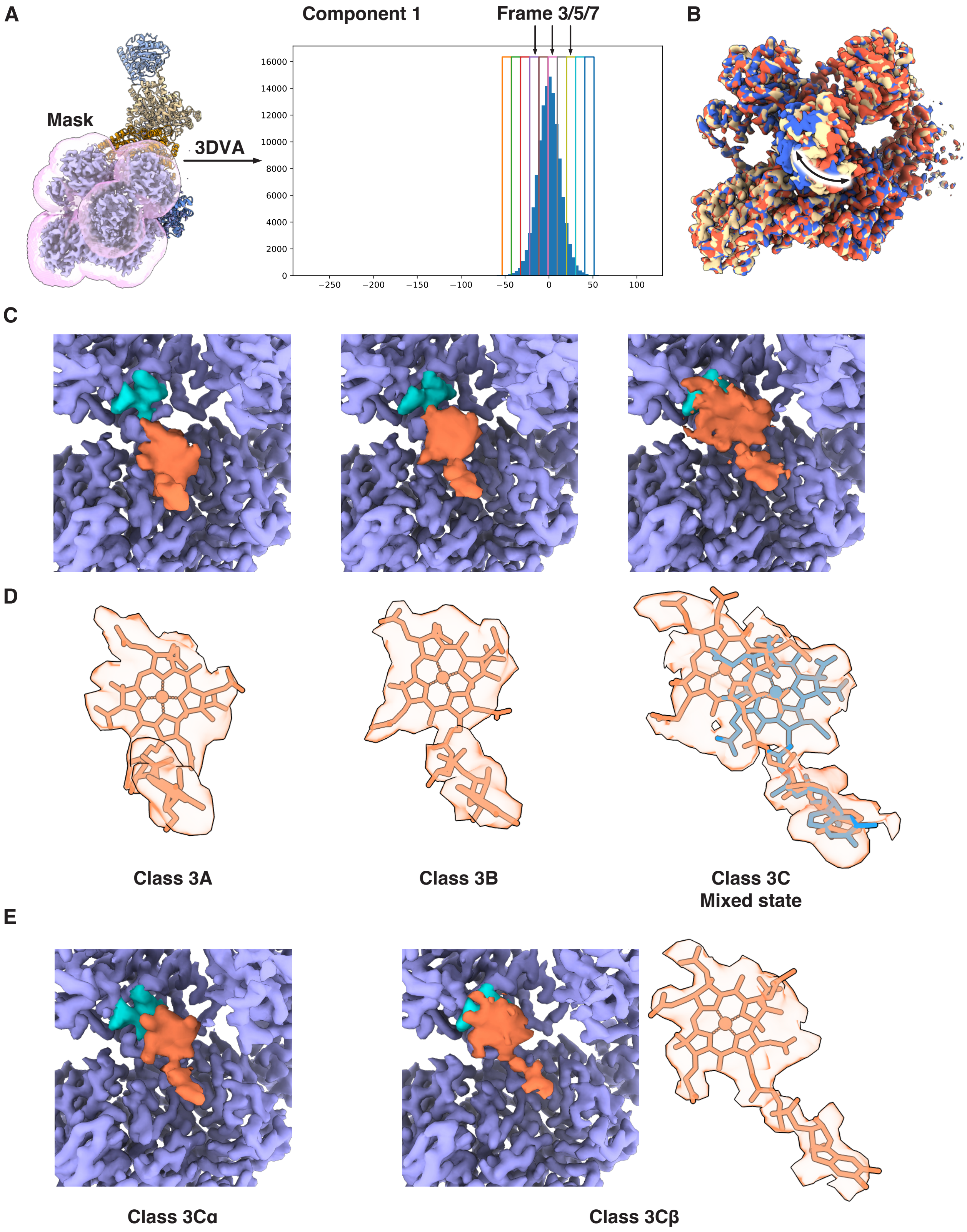
A
Mask
3DVA
Component 1
Frame 3/5/7
16000
14000
12000
10000
8000
6000
4000
2000
0
−250
−200
−150
−100
−50
0
50
100
B
C
D
Class 3A
Class 3B
Class 3C
Mixed state
E
Class 3Cα
Class 3Cβ

**Fig. S19. Motion of the CoFeSP analyzed by 3D variability analysis. (A)** An overview of the CODH/ACS-CoFeSP complex, with the masked region for 3DVA depicted as a light pink surface. Within component 1 output by 3DVA, frames used as references for subsequent 3D focused classification are indicated with vertical arrows. **(B)** Superposition of refined local maps from class 3A (blue), 3B (light yellow) and 3C (red) reveals a rotational movement of the B12 domain. **(C)** Close-up views of these local maps illustrate the approach of B12 toward the A-cluster. Map surfaces are shown in purple, with the A-cluster highlighted in turquoise and B12 in dark orange. **(D)** The map-model fit of B12, with the cryo-EM map density represented as light orange surfaces. The B12 model is depicted in either light orange or light blue. The extended density in class 3C suggests an alternative position of B12 nearer to the A-cluster. **(E)** Close-up views of local maps from class 3Cα and 3Cβ illustrate the further approach of B12 towards the A-cluster. Same color scheme is applied as in (C) and (D).

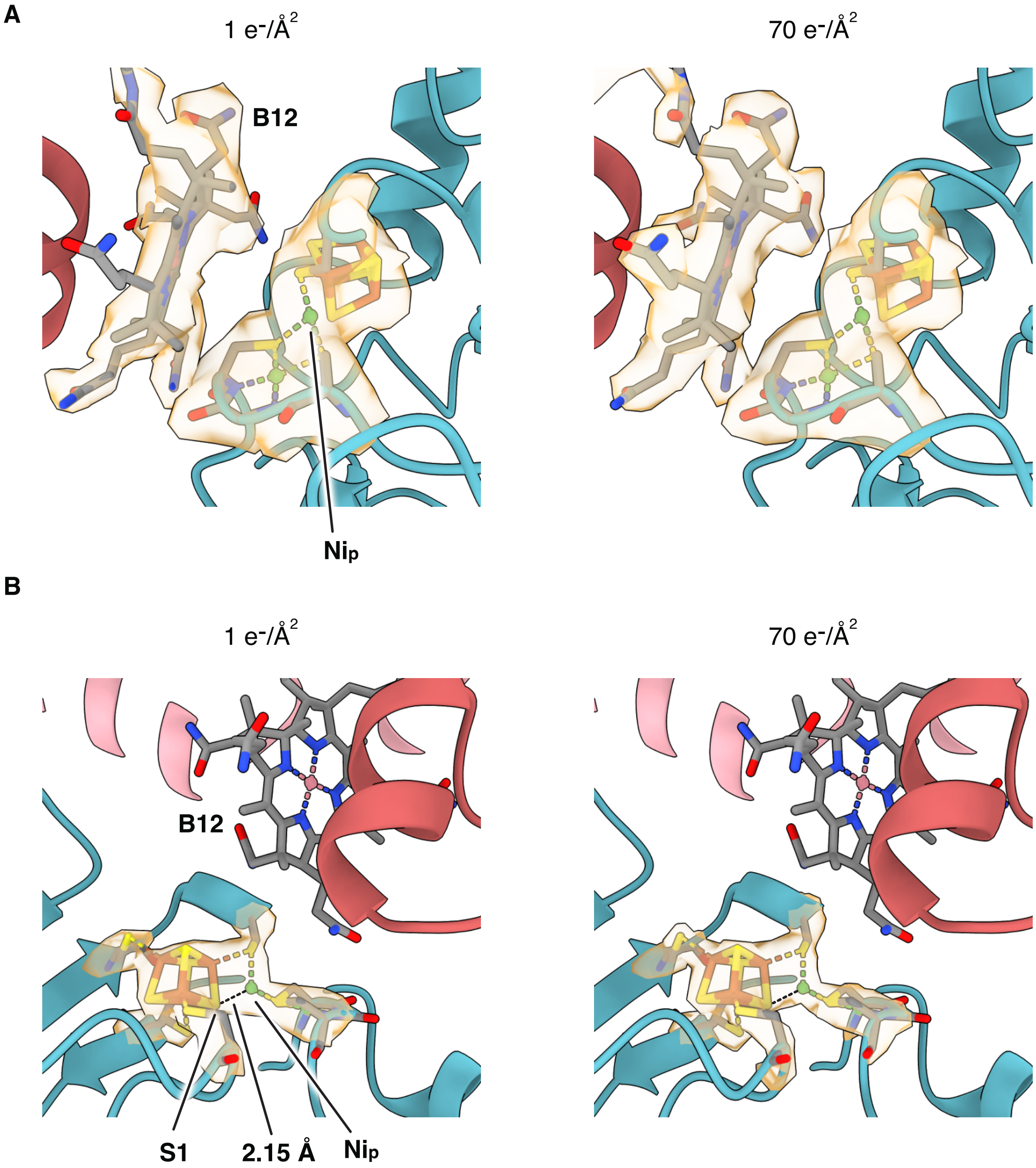


**Fig. S20. Cryo-EM map density of the B12 and A-cluster, reconstructed with various total electron doses. (A)** Close-up views of maps reconstructed from the class 3Cβ reveal that no additional density for a methyl or carbonyl group could be observed at the B12 or the $Ni_p$ site. **(B)** Close-up views of maps reconstructed from the class 3B reveal the proximity of the $Ni_p$ to the S1 of the A-cluster, with distance between $Ni_p$ and S1 indicated by black dashed lines. ACS and CoFeSP are depicted in cartoons, and the B12 and A-cluster in sticks, with their cryo-EM densities shown as light orange surfaces and displayed at 2.58 sigma **(A, left)**, 7.12 sigma **(A, right)**, 5.73 sigma **(B, left)**, and 15.12 sigma **(B, right)**, respectively.

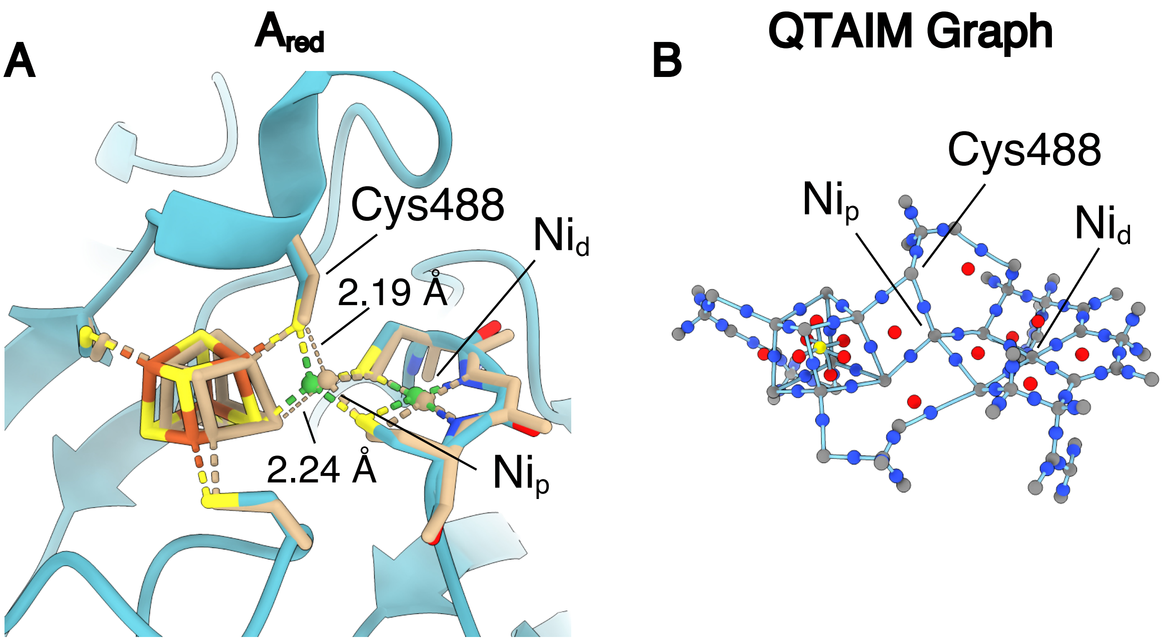


**Fig. S21. Geometry-optimized structure and QTAIM analysis of the A-cluster. (A)** The predicted structure of the A-cluster after geometry optimization with the TPSS-D3(BJ)/def2-TZVP level of theory (superimposed tan structure). Formal oxidation states according to the $A_{red}$ intermediate were used during the calculation. Dotted lines show the distance between $Ni_p$ and the sulfur atoms in the [4Fe-4S] cluster and the coordinating Cys488. The experimental structure is shown in blue (backbone) with Fe in orange, Ni in green, and S in yellow. **(B)** The QTAIM molecular graph based on the topology of the electron density. Critical points are colored according to their classification: nuclear in gray, bond in blue, ring in red, and cage in yellow. Calculated bond paths are represented as continuous light blue lines.

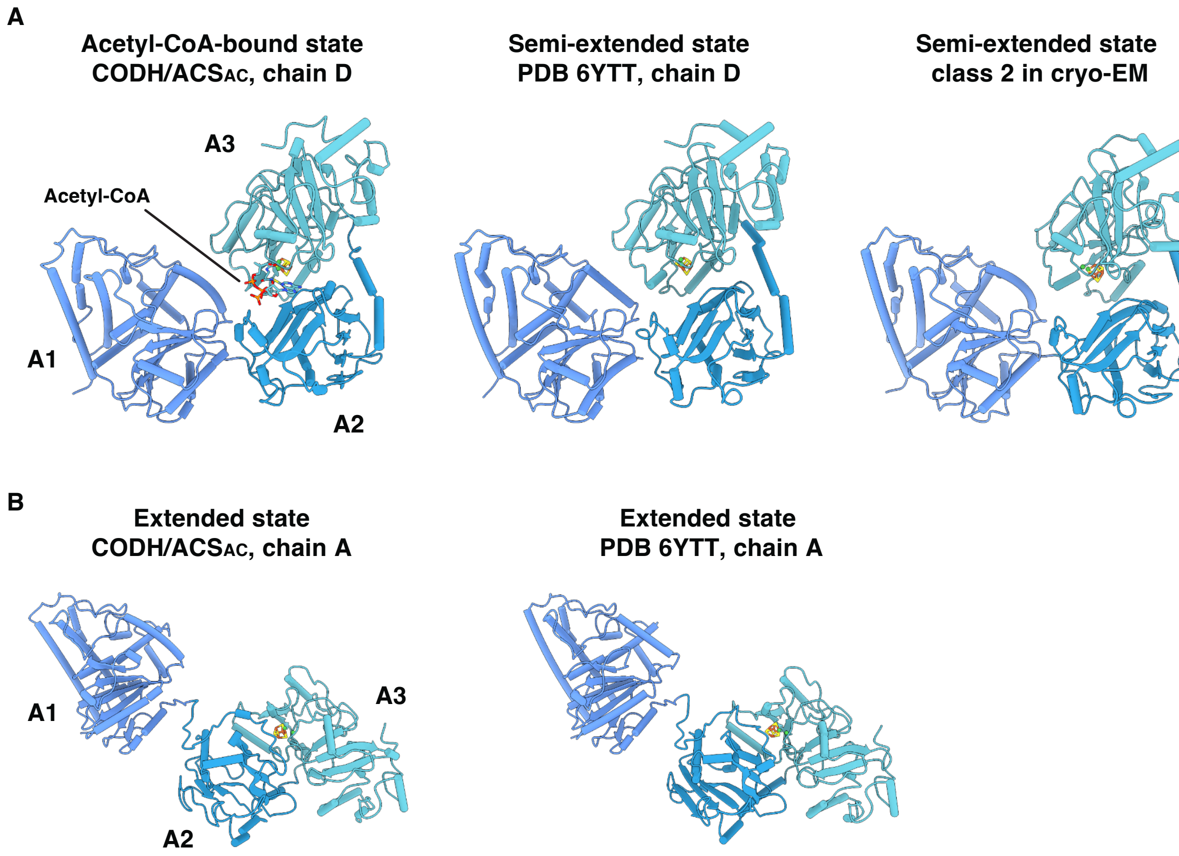


**Fig. S22. Resemblance of CODH/ACS$_{AC}$ conformations to known ACS structures. (A)** Chain D of CODH/ACS$_{AC}$ (acetyl-CoA bound) resembles semi-extended ACSs. **(B)** Chain A of CODH/ACS$_{AC}$ (acetyl-CoA free) exhibits an extended conformation.

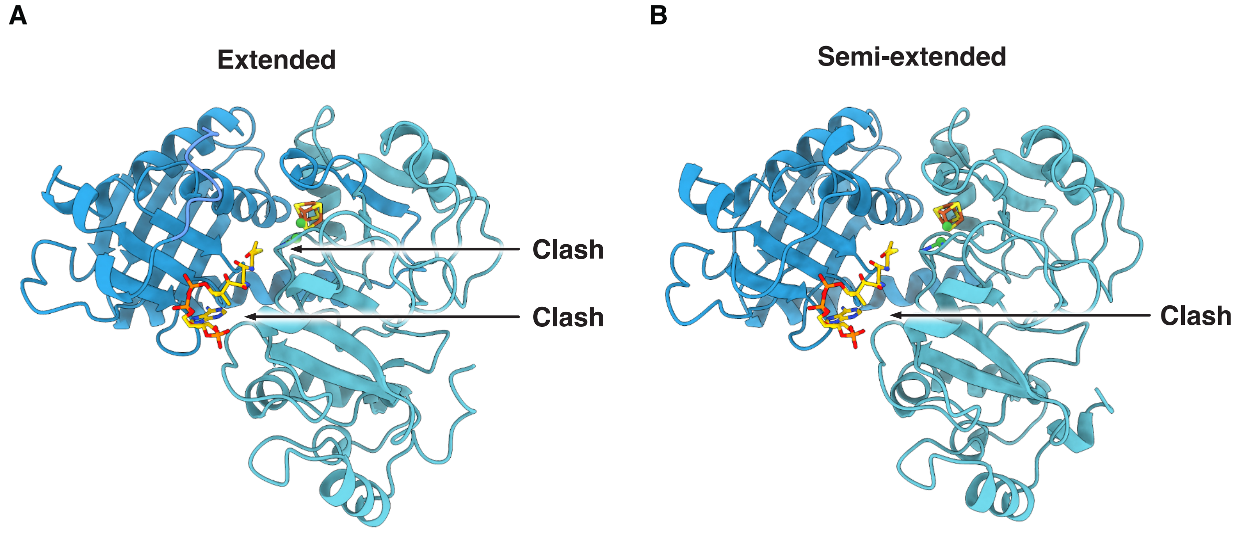


**Fig. S23. Compatibility of the observed acetyl-CoA binding pose with various ACS conformations.** Acetyl-CoA is positioned relative to A2 as observed in CODH/ACS$_{AC}$ (chain D). Clashes (black arrows) occur between acetyl-CoA and A3 in states with a small distance $L_{2-3}$, namely the extended state (PDB 6YTT, chain A (*1*)) **(A)** and the semi-extended state (class 2 in cryo-EM) **(B)**.

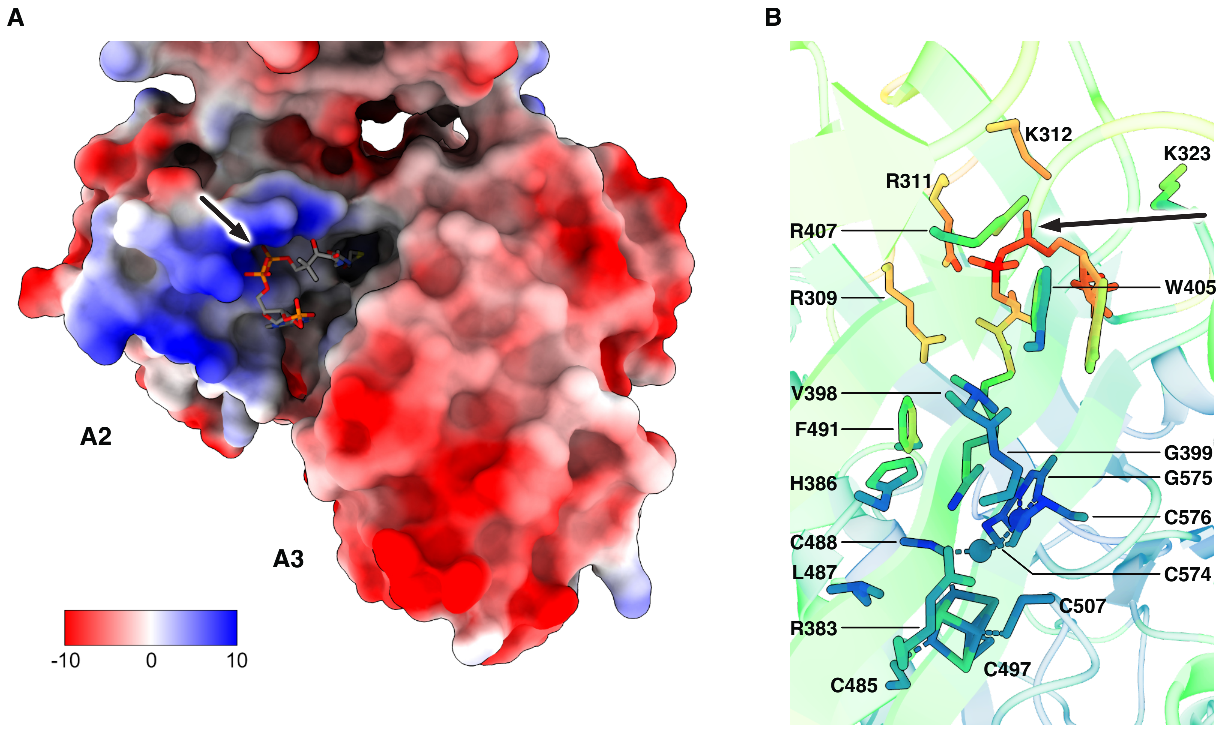


**Fig. S24. The diphosphate moiety of acetyl-CoA faces a positively charged patch on the A2.** The molecular surface of ACS is colored according to the electrostatic potential (in units of kcal/(mol•*e*) at 298 K) in ChimeraX (*16*). The acetyl-CoA is depicted as sticks and colored according to element. **(B)** The CODH/ACS$_{AC}$ colored by b-factors, viewed from the same angle as in Fig. 4B. Red indicates high b-factors, while dark blue indicates low b-factors. The diphosphate moiety of acetyl-CoA is marked by arrows in both panels.

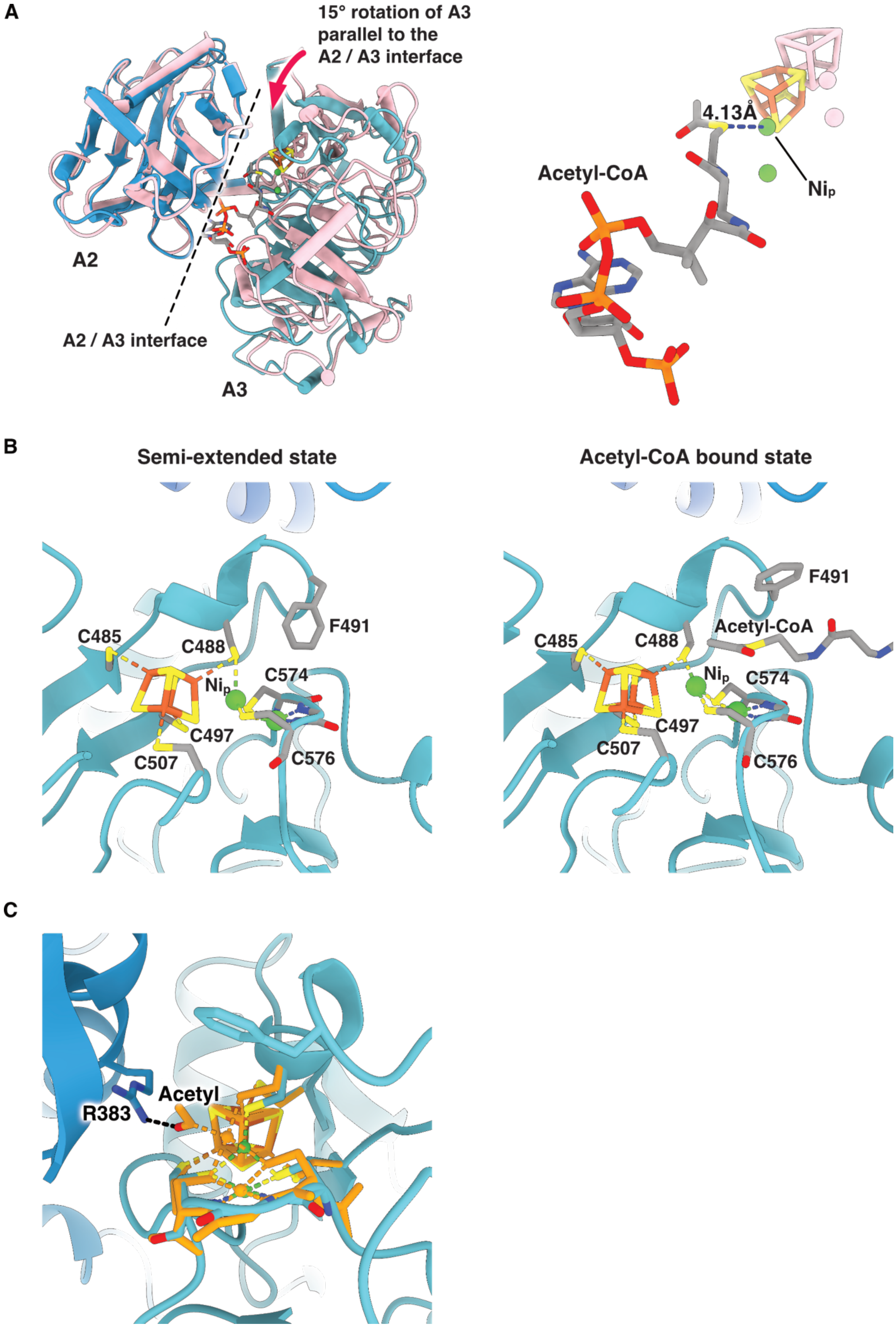
A
15° rotation of A3 parallel to the A2 / A3 interface
A2
A2 / A3 interface
A3
4.13Å
Acetyl-CoA
Nip
B
Semi-extended state
Acetyl-CoA bound state
C485
C488
F491
Nip
C574
C497
C507
C576
C485
C488
F491
Acetyl-CoA
Nip
C574
C497
C507
C576
C
Acetyl
R383

**Fig. S25. Fine tuning of the ACS conformation for acetylation of the CoA. (A)** A 15° rotation of A3 (left panel) brings the A-cluster into close proximity to CoA or acetyl-CoA (right panel), revealed by alignment of the acetyl-CoA-bound ACS (CODH/$ACS_{AC}$, chain D; blue cartoon) and a semi-extended ACS (class 2 in cryo-EM; pink cartoon) on the basis of the A2 domain. Distance measurement between the $Ni_p$ and the sulfur atom of acetyl-CoA is indicated by a dashed line. For clarity, only A2 and A3 are visualized. **(B)** Displacement of Phe491 for CoA or acetyl-CoA binding at the A-cluster. Compared to the semi-extended state (PDB 6YTT (*1*), chain D), the acetyl-CoA-bound state (CODH/$ACS_{AC}$, chain D) shows movement of F491 away from the A-cluster to accommodate acetyl-CoA and avoid steric clashes. The equivalent residue to F491 is F512 in *Mt*ACS. **(C)** Arg383 may stabilize a $Ni_p$-bound acetyl by hydrogen bonding. A DFT-optimized structure of the acetyl-bound A-cluster ($Ni_p$(III)-Ac state) (*36*) is superimposed upon our acetyl-CoA-bound structure. Acetyl-CoA in the CODH/$ACS_{AC}$ is omitted for clarity. In this conformation, Arg383 of A2 is well-positioned to stabilize a $Ni_p$-bound acetyl group by hydrogen bonding (dashed line). The DFT-optimized structure is colored in orange, except for the oxygen atom of the acetyl group, which is in red.

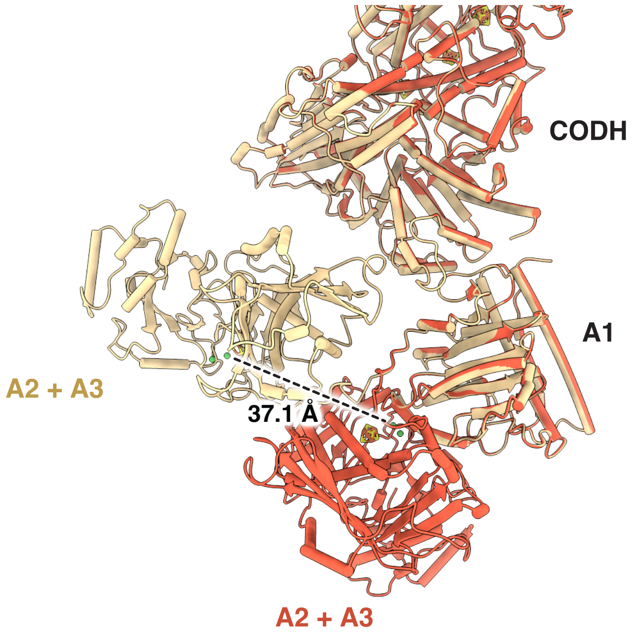


**Fig. S26. Between the closed and hyper-extended states, the A-cluster undergoes a displacement of over 37 Å.** Comparison of the closed ACS (class 1 in cryo-EM, red cartoon) with a hyper-extended ACS (class 3Cβ in cryo-EM, light-yellow cartoon) superimposed according to A1. The distance measurement between the $Ni_p$ in both states is marked by a dashed line.

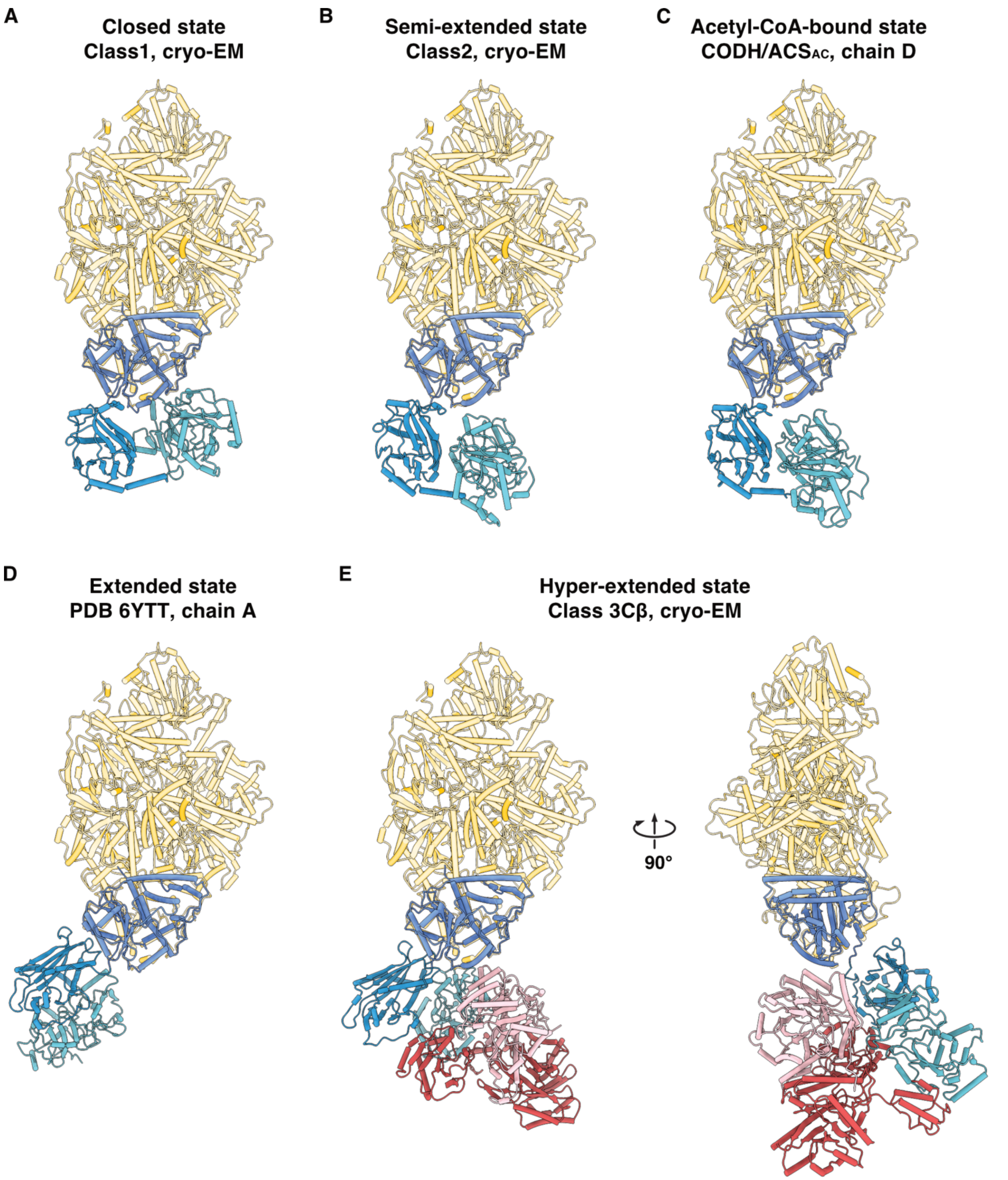


**Fig. S27. All identified *Ca*ACS conformations are compatible with the overall architecture of *Mt*CODH/ACS. (A-E)** Each conformation of *Ca*ACS is aligned with a *Mt*CODH/ACS (PDB 1OAO (*47*), chain D; yellow) according to the A1 domain. No clashes are observed between the *Mt*CODH/ACS rigid core and *Ca*ACS A2 or A3 domain. **(E)** *Ca*CoFeSP in complex with hyper-extended *Ca*ACS also exhibits no clashes. For clarity, only *Ca*ACS, *Ca*CoFeSP and the rigid core of *Mt*CODH/ACS are displayed.

**Table S1. EM statistics**

| | |
|---|---|
| Data collection | |
| Electron microscope | Titan Krios G4 |
| Camera | Falcon 4 (electron-counting mode) |
| Data collection software | EPU with aberration-free image shift (AFIS) |
| Voltage | 300 kV |
| Nominal magnification | 165,000 x |
| Calibrated pixel size | 0.73 Å/pix |
| Dose rate | 5.02 / 4.46 / 4.63 $e^{-}$/pix/s* |
| Total exposure | 70 $e^{-}Å^{-2}$ |
| Number of frames | 1946 / 2191 / 2114 raw frames† |
| Defocus range | -0.8 to -2.5 µm |
| Energy filter slit width | 8.0 eV |
| Image processing | |
| Motion correction software | MotionCor2 (RELION implementation) |
| CTF estimation | CTFFIND 4.1.13 |
| Particle selection software | Topaz |
| Exposures used | 39,159 movies in EER format |
| Particles extracted | 5,819,001 |
| Particles contributing to final datasets | 1,873,249‡ |
| Classification software | RELION 4 and CryoSPARC 4 |
| Refinement software | CryoSPARC 4 |
| Model building | |
| Modeling software | Coot |
| Refinement software | PHENIX real-space refinement |

* Respective dose rates for three datasets.
† Respective counts of raw frames for three datasets.
‡ Dimeric particles; effective particle number is doubled in processing by $C_2$ symmetry expansion.

### Table S2. Cryo-EM map identifiers and quality statistics

| EMDB ID | Description | Resolution in Å ($FSC_{0.143}$) | Sharpening B-factor | # of particles | Symm. |
|---|---|---|---|---|---|
| EMD-50898 | CODH/ACS rigid core (dataset 1) | 1.94 | -46.9 | 649,146 | $C_2$ |
| EMD-50899 | CoFeSP-bound ACS in class 3A | 2.71 | -60.7 | 140,666* | $C_1$ |
| EMD-50901 | CoFeSP-bound ACS in class 3B | 2.65 | -57.6 | 129,529* | $C_1$ |
| EMD-50903 | CoFeSP-bound ACS in class 3Cβ | 2.88 | -57.0 | 76,451* | $C_1$ |
| EMD-50904 | CoFeSP-bound ACS in class 3Cα | 2.78 | -56.1 | 72,324* | $C_1$ |
| EMD-50906 | ACS in closed and CO-bound state | 2.83 | -59.5 | 255,954* | $C_1$ |
| EMD-50908 | ACS in semi-extended state | 3.29 | -37.6 | 52,916* | $C_1$ |
| EMD-50909 | CODH/ACS in ferredoxin-bound state, locally filtered map | 2.10 | n/a | 896,132 | $C_1$ |
| EMD-50897 | CoFeSP-bound CODH/ACS in class 3A, generated from EMD-50898 and EMD-50899 | Composite map | n/a | n/a | n/a |
| EMD-50900 | CoFeSP-bound CODH/ACS in class 3B, generated from EMD-50898 and EMD-50901 | Composite map | n/a | n/a | n/a |
| EMD-50902 | CoFeSP-bound CODH/ACS in class 3Cβ, generated from EMD-50898 and EMD-50903 | Composite map | n/a | n/a | n/a |
| EMD-50905 | CODH/ACS with ACS in closed and CO-bound state, generated from EMD-50898 and EMD-50906 | Composite map | n/a | n/a | n/a |
| EMD-50907 | CODH/ACS with ACS in semi-extended state, generated from EMD-50898 and EMD-50908 | Composite map | n/a | n/a | n/a |

*Particle counts after $C_2$ symmetry expansion.

**Table S3. Broken Symmetry-DFT energetics of the C-cluster.** Energies of the broken symmetry states explored for the different oxidation states of the C-cluster, using the TPSS-D3(BJ)/def2-TZVP level of theory. Energies of the antiferromagnetic alignments are presented relative to the ferromagnetic, high-spin state.

| $Fe_a$ | $Fe_b$ | $Fe_c$ | $Fe_d$ | Relative energy [Oxidized] (eV) | Relative energy [$C_{red1(-OH)}$] (eV) | Relative energy [$C_{red2(-OH)}$] (eV) |
|---|---|---|---|---|---|---|
| ↑ | ↑ | ↑ | ↑ | 0.000 | 0.000 | 0.000 |
| ↑ | ↑ | ↓ | ↓ | -1.674 | -0.829 | -0.160 |
| ↑ | ↓ | ↑ | ↓ | -1.649 | -1.465 | -0.366 |
| ↑ | ↓ | ↓ | ↑ | -1.987 | -1.366 | -0.211 |
| ↓ | ↑ | ↑ | ↓ | -1.973 | -1.115 | -0.317 |
| ↓ | ↑ | ↓ | ↑ | -1.887 | -1.140 | -0.356 |
| ↓ | ↓ | ↑ | ↑ | -1.422 | -1.236 | -0.144 |

**Table S4. Model identifiers and quality statistics**

| | |
|---|---|
| CoFeSP-bound class 3A | |
| PDB ID | 9FZY |
| EMDB ID | EMD-50897 |
| Residues built | 3009 |
| Ligands built | 5x SF4, 1x B12, 2x NI, 2x RQM |
| RMS bond lengths | 0.006 |
| RMS bond angles | 1.091 |
| Ramachandran outliers | 0.07% |
| Ramachandran favored | 96.96% |
| Rotamer outliers | 0.73% |
| Clashscore | 4.11 |
| CaBLAM outliers | 0.97% |
| C-beta outliers | 0.00% |
| CoFeSP-bound class 3B | |
| PDB ID | 9FZZ |
| EMDB ID | EMD-50900 |
| Residues built | 3009 |
| Ligands built | 5x SF4, 1x B12, 2x NI, 2x RQM |
| RMS bond lengths | 0.008 |
| RMS bond angles | 1.116 |
| Ramachandran outliers | 0.10% |
| Ramachandran favored | 97.30% |
| Rotamer outliers | 0.52% |
| Clashscore | 3.81 |
| CaBLAM outliers | 0.90% |
| C-beta outliers | 0.00% |
| CoFeSP-bound class 3Cβ | |
| PDB ID | 9G00 |
| EMDB ID | EMD-50902 |
| Residues built | 3009 |
| Ligands built | 5x SF4, 1x B12, 2x NI, 2x RQM |
| RMS bond lengths | 0.007 |
| RMS bond angles | 1.096 |
| Ramachandran outliers | 0.07% |
| Ramachandran favored | 97.26% |
| Rotamer outliers | 0.73% |
| Clashscore | 3.33 |
| CaBLAM outliers | 0.84% |
| C-beta outliers | 0.00% |

| CODH/ACS with ACS in closed / CO-bound state | |
|---|---|
| PDB ID | 9G01 |
| EMDB ID | EMD-50905 |
| Residues built | 2254 |
| Ligands built | 4x SF4, 2x NI, 2x RQM, 1x CMO |
| RMS bond lengths | 0.009 |
| RMS bond angles | 1.116 |
| Ramachandran outliers | 0.09% |
| Ramachandran favored | 97.33% |
| Rotamer outliers | 0.54% |
| Clashscore | 2.82 |
| CaBLAM outliers | 0.98% |
| C-beta outliers | 0.00% |
| **CODH/ACS with ACS in semi-extended state** | |
| PDB ID | 9G02 |
| EMDB ID | EMD-50907 |
| Residues built | 2253 |
| Ligands built | 4x SF4, 2x NI, 2x RQM |
| RMS bond lengths | 0.008 |
| RMS bond angles | 1.115 |
| Ramachandran outliers | 0.09% |
| Ramachandran favored | 96.93% |
| Rotamer outliers | 0.97% |
| Clashscore | 4.40 |
| CaBLAM outliers | 0.98% |
| C-beta outliers | 0.00% |
| **CODH/ACS in ferredoxin-bound state** | |
| PDB ID | 9G03 |
| EMDB ID | EMD-50909 |
| Residues built | 1890 |
| Ligands built | 5x SF4, 2x RQM |
| RMS bond lengths | 0.005 |
| RMS bond angles | 1.079 |
| Ramachandran outliers | 0.11% |
| Ramachandran favored | 97.45% |
| Rotamer outliers | 0.39% |
| Clashscore | 2.34 |
| CaBLAM outliers | 0.91% |
| C-beta outliers | 0.00% |

**Table S5. RMSDs of ACS domains aligned with an *Mt*ACS reference structure.** To assess the rigidity of ACS domains, individual domains from various structures were aligned with an *Mt*ACS reference structure (PDB 1MJG (*48*) chain M) using ChimeraX's matchmaker tool (*16*). The RMSDs in Å over all residue pairs, as reported by the matchmaker, are presented in the table.

| RMSD vs. 1MJG chain M | A1* | A2 † | A3‡ |
|---|---|---|---|
| 1MJG chain M | - | - | - |
| 1OAO chain C | 0.194 | 0.406 | 0.341 |
| 1OAO chain D | 0.636 | 0.712 | 0.399 |
| 1RU3 | 0.854 | 0.730 | 0.587 |
| 6X5K chain M | 0.132 | 0.183 | 0.173 |
| 6YTT chain A | 1.634 | 1.047 | 0.751 |
| 6YTT chain D | 1.663 | 1.019 | 0.727 |
| 7ZKJ chain B | 0.398 | 0.666 | 0.567 |
| CryoEM closed (9G01) | 1.609 | 0.903 | 0.939 |
| CryoEM semi-extended (9G02) | 1.618 | 1.062 | 1.003 |
| CryoEM class 3A (9FZY) | 1.630 | 0.905 | 0.843 |
| CryoEM class 3B (9FZZ) | 1.629 | 0.973 | 0.801 |
| CryoEM class 3Cβ (9G00) | 1.629 | 0.992 | 0.811 |
| 9G7I chain D | 1.747 | 1.075 | 1.097 |
| 9G7I chain A | 1.649 | 1.609 | 0.956 |

* A1 definition in current study: *Mt*ACS 20-309, *Ca*ACS 2-286, *Ch*ACS 23-312;
† A2 definition in current study: *Mt*ACS 326-494, *Ca*ACS 303-473, *Ch*ACS 329-497;
‡ A3 definition in current study: *Mt*ACS 502-729, *Ca*ACS 481-708, *Ch*ACS 505-732.

**Table S6. Interdomain distances within ACSs.** Distance between A1 and A3 is represented by the distance between F209 (Cα) and $Ni_p$ (more broadly, $M_p$, due to variations in metal binding across structures); distance between A2 and A3 is indicated by the distance between F491 (Cα) and W405 (Cα).

| | F209-$M_p$ (Å)* | F491-W405 (Å)† |
|---|---|---|
| 1MJG chain M | 7.478 | 34.472 |
| 1OAO chain C | 6.731 | 34.344 |
| 1OAO chain D | 11.788 | 35.206 |
| 1RU3 | 13.562 | 31.657 |
| 6X5K chain M | 7.000 | 34.277 |
| 6YTT chain A | 44.890 | 20.400 |
| 6YTT chain D | 25.347 | 17.468 |
| 7ZKJ chain B | 7.146 | 33.499 |
| CryoEM closed (9G01) | 7.084 | 35.153 |
| CryoEM semi-extended (9G02) | 25.405 | 20.442 |
| CryoEM class 3A (9FZY) | 39.823 | 36.784 |
| CryoEM class 3B (9FZZ) | 39.794 | 37.842 |
| CryoEM class 3Cβ (9G00) | 40.490 | 39.350 |
| 9G7I chain D | 26.652 | 14.852 |
| 9G7I chain A | 44.447 | 20.313 |

* F209 is equivalent to F229 in *Mt*ACS and F232 in *Ch*ACS;
† F491 is equivalent to F512 in *Mt*ACS and F515 in *Ch*ACS; W405 is equivalent to W427 in *Mt*ACS and W430 in *Ch*ACS.

**Table S7. Broken Symmetry-DFT energetics of the A-cluster.** Energies of the broken symmetry states of the $[4Fe\text{-}4S]^{2+}$ cubane within the A-cluster with TPSS-D3(BJ)/def2-TZVP level of theory. All values are relative to the energy of the fully ferromagnetic, high-spin state.

| $Fe_a$ | $Fe_b$ | $Fe_c$ | $Fe_d$ | Relative energy (eV) |
|---|---|---|---|---|
| ↑ | ↑ | ↑ | ↑ | 0.000 |
| ↑ | ↑ | ↓ | ↓ | -2.142 |
| ↑ | ↓ | ↑ | ↓ | -2.129 |
| ↑ | ↓ | ↓ | ↑ | -2.134 |
| ↓ | ↑ | ↑ | ↓ | -2.140 |
| ↓ | ↑ | ↓ | ↑ | -1.795 |
| ↓ | ↓ | ↑ | ↑ | -1.997 |

**Table S8. X-ray analysis statistics for the crystallographic structure.**

| | CODH/ACS in complex with Acetyl-CoA |
|---|---|
| **Data collection** | |
| Synchrotron source | SLS, X06DA |
| Wavelength (Å) | 1.00003 |
| Space group | $P4_22_12$ |
| Resolution (Å) | 59.61 − 2.93 (3.16 −2.93) |
| Cell dimensions | |
| a, b, c (Å) | 298.08, 298.08, 127.53 |
| α, β, γ (°) | 90, 90, 90 |
| $R_{merge}$(%)* | 20.7 (242.3) |
| $R_{pim}$ (%)* | 4.5 (51.3) |
| $CC_{1/2}$* | 0.998 (0.593) |
| $I/\sigma_I$* | 14.5 (1.6) |
| Spherical completeness* | 81.4 (21.0) |
| Ellipsoidal completeness* | 96.1 (73.0) |
| Redundancy* | 22.1 (23.3) |
| Nr. unique reflections* | 99,666 (4,983) |
| **Refinement** | |
| Resolution (Å) | 49.68 − 2.93 |
| Number of reflections | 99,636 |
| $R_{work}/R_{free}$ † (%) | 19.02/22.02 |
| Number of atoms | |
| Protein | 20,141 |
| Solvent and ligands | 342 |
| Water | 14 |
| Mean B-value ($Å^2$) | 89.64 |
| Molprobity clash score, all atoms | 3.75 |
| Ramachandran plot | |
| Favored regions (%) | 94.86 |
| Outlier regions (%) | 0.19 |
| RMSD bond lengths (Å) | 0.010 |
| RMSD bond angles (°) | 1.367 |
| **PDB ID code** | **9G7I** |

* Values relative to the highest resolution shell are within parentheses.
† $R_{free}$ was calculated as the $R_{work}$ for 5 % of the reflections that were not included in the refinement.

**Movie S1.**

The conformation-based model of acetyl-CoA synthesis, involving initial electron transfer from ferredoxin, $CO_2$ reduction, and conformational changes of the ACS to facilitate carbonylation and methylation of the A-cluster, culminating in the acetylation of CoA.
